# Lysosomal Expansion Compartments Mediate Zinc and Copper Homeostasis in *Caenorhabditis elegans*

**DOI:** 10.64898/2026.03.05.709934

**Authors:** Joy R. Armendariz, Sean Teng, Camdyn Rakow, Raquel Herrera, Samuel Herrera, Moshe T. Gordon, Si Chen, Stefan Vogt, Hanwenheng Liu, Makena Jarvis, Katryna Reese, Aidan T. Pezacki, Christopher J. Chang, Byung-Eun Kim, Daniel L. Schneider, Adelita D. Mendoza, Kerry Kornfeld

**Affiliations:** Department of Biochemistry, University of Colorado, Boulder, Colorado, 80303; Department of Developmental Biology, Washington University School of Medicine, Saint Louis, Missouri, 63110; X-ray Science Division, Advanced Photon Source, Argonne National Laboratory, Lemont, Illinois, 60439; Department of Chemistry, Princeton University, Princeton, New Jersey 08544; Department of Chemistry, University of California at Berkeley, Berkeley, California 94720; Department of Animal and Avian Sciences, University of Maryland, College Park, Maryland 20742

## Abstract

Zinc is an essential transition metal that participates in many biological processes. In *C. elegans*, excess zinc is stored in lysosomes in intestinal cells; this process involves increasing the expression of the zinc transporter CDF-2 and remodeling of lysosomes characterized by an increase in the volume of the expansion compartment. To determine if this is a more general property, we investigated other metals. Here we report that lysosomes are remodeled in response to excess copper, manganese, and cadmium, with each metal causing an increase in the volume of the expansion compartment. Mutants with a reduced number of lysosomes were hypersensitive to growth retardation caused by excess copper and manganese, suggesting metal toxicity is prevented by metal sequestration in lysosomes. Using a novel method to analyze isolated lysosomes by X-ray Fluorescence Microscopy we demonstrated that zinc, copper and manganese are detectable in the lumen of lysosomes. To further analyze copper, we examined localization of CUA-1.1, a copper transporter that moves copper into the lumen of lysosomes. Like the zinc transporter CDF-2, CUA-1.1 localizes to both the acidified and expansion compartments in excess copper. These results indicate that the same intestinal lysosomes store zinc, copper and manganese. Lysosome remodeling characterized by an increase in volume of the expansion compartment is not specific to zinc but is a more general phenomenon during metal storage in lysosomes.

## Introduction

Zinc is a transition metal that all organisms require for survival.^1,2^ Zinc is predicted to bind approximately 10% of the human proteome, corresponding to ∼3000 proteins.^3^ Protein bound zinc plays essential roles in enzyme catalysis_3_ and protein structure^4^, whereas labile zinc functions as a signal transduction molecule.^5^ Given this widespread requirement, organisms have evolved sophisticated zinc homeostasis mechanisms to tightly regulate intracellular zinc levels. Dysregulation of zinc results in either zinc deficiency or zinc toxicity and can lead to human pathologies such as acrodermatitis enteropathica^6^ and neurodegeneration.^7,8^ Zinc homeostasis is partly mediated by transmembrane proteins that transport zinc across membranes. SLC30A/ZNT/CDF family members transport zinc from the cytosol to extracellular spaces or luminal spaces within cells._9,10_ Members of the SLC39A/Zrt,Irt-like (ZIP) family proteins transport zinc into the cytosol from intracellular compartments or extracellular spaces.^9,10^

*Caenorhabditis elegans* has been used to study zinc homeostasis because this tractable model organism offers powerful genetic analysis, live fluorescence imaging enabled by a translucent body, and conservation with humans (approximately 38% of *C. elegans* genes are orthologous to human genes).^11^ In *C. elegans*, high- and low- zinc homeostasis pathways have been identified.^12,13^ In excess zinc conditions, the high zinc sensor HIZR-1, a nuclear receptor transcription factor, senses elevated cytosolic zinc levels by binding zinc at its ligand-binding domain. Zinc bound HIZR-1 translocates to the nucleus, where the DNA-binding domain interacts with high specificity and affinity with the High Zinc Activation (HZA) enhancer element.^14,15^ This leads to the upregulation of multiple genes that are active during high zinc homeostasis, including *cdf-2*. The CDF-2 protein transports zinc from the cytosol into the lumen of intestinal lysosomes, resulting in zinc storage and detoxification.^16,17^ The low-zinc homeostasis pathway is less well defined but involves the activation of the Low Zinc Activation (LZA) enhancer element, resulting in the upregulation of low-zinc homeostasis genes, including *zipt-2.3*.^13^ The ZIPT-2.3 protein transports zinc from the lumen of lysosomes into the cytosol.^18^ The CDF-2 and ZIPT-2.3 protein are reciprocally regulated by zinc levels at the transcriptional, protein, and physiological levels.^18^

Lysosomes in intestinal cells of *C. elegans* contain two compartments – an acidified compartment, and a dynamic expansion compartment that does not appear to be acidic. Changes in zinc concentration lead to morphological restructuring in lysosomes.^18,17^ In zinc-replete conditions, the acidified compartment predominates, whereas the expansion compartment is small. In zinc-deficient conditions, the expansion compartment is moderately increased in volume, while in zinc-excess conditions, the expansion compartment is dramatically increased in volume.^18^ In each case, CDF-2 localizes to the membrane of the acidified compartment and the expansion compartment, whereas ZIPT-2.3 has been shown to localize to the membrane of the acidified compartment but not the expansion compartment.^18^ This morphological remodeling is predicted to facilitate a change in the ratio of CDF-2 and ZIPT-2.3 on the membrane of the lysosome, resulting in a net flow of zinc into lysosomes in zinc-excess conditions or a net flow of zinc into the cytosol in zinc-deficient conditions.

The findings that zinc is stored in lysosomes and excess-zinc induces morphological remodeling led us to ask if other transition metals are also stored in lysosomes and induce lysosome remodeling. Copper and manganese are essential metals necessary for a wide range of cellular processes such as cell fate, mitochondrial function, immune function, digestion, and metabolism.^19–21^ Cadmium is not an essential metal but it is a potent toxin – it is the same group as zinc on the periodic table and is predicted to displace zinc from important binding sites in proteins.^22–24^ Like zinc, copper also requires tight homeostatic regulation; dysregulation of copper leads to oxidative cellular stress^25,26^ and human pathologies such as Wilson’s and Menkes diseases.^27,28^ Lysosomes have been linked to copper homeostasis and are emerging as an important site for copper trafficking. In *C. elegans*, intestinal cells are a key site for copper homeostasis; in replete copper conditions, the copper transporter CUA-1.1 (ATP7A/B) localizes to the basolateral membrane where it transports copper into the pseudocoelom.^29^ However, exposure to excess copper causes re-localization of CUA-1.1 to the membrane of lysosomes where it transports copper into the lysosomal lumen.^29^ This contrasts with the zinc transporter CDF-2, which constitutively localizes to lysosomes and is not known to function elsewhere. Regulation of *cua-1.1* also contrasts with *cdf-2*, because exposure to copper leads to downregulation of *cua-1.1* mRNA, while exposure to zinc leads to upregulation of *cdf-2* mRNA.^16,29^ Although CUA-1.1 localizes to lysosomes in excess copper, its relationship to the expansion compartment remains unknown.

Here we report that lysosomes are remodeled in response to excess copper, manganese, and cadmium, with each condition causing an increase in the volume of the expansion compartment. Mutants with a reduced number of lysosomes were hypersensitive to growth retardation caused by excess copper and manganese, suggesting these metals are detoxified by sequestration in lysosomes. To quantify the metal content of intestinal lysosomes, we developed a new method to analyze isolated lysosomes by X-ray Fluorescence Microscopy (XFM). Zinc, copper and manganese were detectable in the lumen of lysosomes. To probe the mechanism of copper storage, we analyzed the localization of CUA-1.1; like the zinc transporter CDF-2, the copper transporter CUA-1.1 localizes to both the acidified and expansion compartments in excess copper. These results indicate that the same intestinal lysosomes may store zinc, copper and manganese, and lysosome remodeling characterized by an increase in volume of the expansion compartment is not specific to zinc but is a more general phenomenon during metal storage in lysosomes.

## Results

### Excess zinc, copper, manganese, and cadmium induce morphological remodeling of lysosomes

Excess zinc causes morphological changes in lysosomes in intestinal cells; remodeled lysosomes are characterized by an increase in the volume of the expansion compartment. To determine if this response is specific for zinc, we analyzed the response to the physiological transition metals copper and manganese and the heavy metal cadmium. L4 stage animals expressing CDF-2::GFP and ZIPT-2.3::mCherry were exposed to 200µM supplemental zinc, 100μM supplemental copper, 20mM supplemental manganese, 200μM cadmium or control not supplemented with any of the proceeding metals for 24 hours on NAMM dishes containing *E. coli* OP50 supplemented with LysoTracker Blue. CDF-2 is a zinc importer that localizes to the expansion compartment and the acidified compartment, whereas ZIPT-2.3 is a zinc exporter that localizes to the acidified compartment but not the expansion compartment.^18^ LysoTracker also localizes to the acidified compartment but not the expansion compartment. Lysosomes from control worms cultured on NAMM displayed a spherical morphology; CDF-2::GFP, ZIPT-2.3::mCherry, and LysoTracker Blue displayed highly overlapping patterns of expression (Figure 1A,B). In excess zinc, the expansion compartment appeared to significantly increase in volume (Figure 1A,C). To quantify this morphological change, we measured the volume of the acidified compartment and the expansion compartment from 3-dimensional super-resolution images. Compared to animals on control medium, animals on excess zinc displayed a significant increase in total volume of the lysosome from ∼4.3 μm_3_ to ∼18.2 μm_3_ (∼4.2X increase) (Figure S1A, Table S1). The volume of the acidified compartment did not change significantly, whereas the volume of the expansion compartment increased significantly from ∼0.73 μm_3_ to ∼15.6 μm_3_ (∼21X increase) (Figure S1C,E). In animals with replete zinc, the expansion compartment represents ∼33% of the total volume of the lysosome - this increased significantly to ∼83% in zinc excess (Figure S1B,D,F). These results are consistent with previous reports.^17,18^

**Figure 1.**
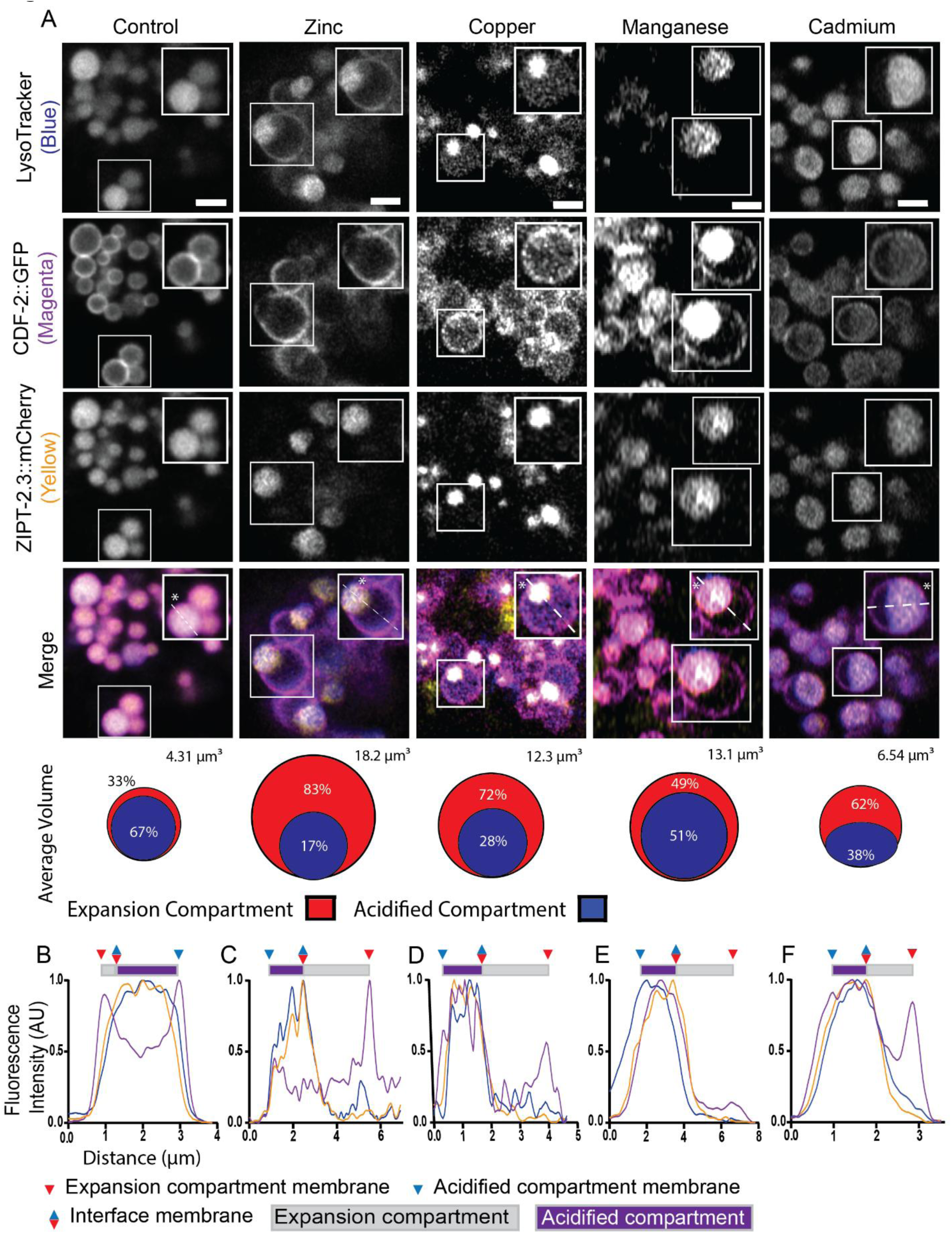
Excess zinc, copper, manganese, and cadmium induce morphological remodeling of lysosomes. (A) Transgenic L4 stage animals expressing CDF-2::GFP (magenta) and ZIPT-2.3::mCherry (yellow) were cultured for 24 h in LysoTracker Blue (blue) on either NAMM medium (control), or NAMM medium containing 200 µM supplemental zinc, 100μM supplemental copper, 20mM supplemental manganese, or 200μM cadmium, and imaged by super-resolution microscopy. Images show a portion of the intestine of 1-day old adult animals. White boxes are enlarged in the upper right of each image; Scale bar = 2 µm. Dashed white lines and asterisks indicate the lysosome analyzed below. Cartoons illustrate the average total volume of a lysosome (μm_3_) and the percent represented by the acidified (blue) and expansion (red) compartments (Figure S1, Table S1). (B-F) Line scans of individual lysosomes in control (B), excess zinc (C), excess copper (D), excess manganese (E), and cadmium (F). Fluorescence intensity along the path of the line was measured beginning at the white asterisk. CDF::GFP, ZIPT-2.3:mCherry, and LysoTracker, are displayed as magenta, yellow, and blue, respectively. For each color line, the highest value of fluorescence intensity is set to 1.0 in arbitrary units (AU), and the lowest fluorescence intensity is set to 0. All other values are normalized. Triangles indicate positions of membranes, and rectangles indicate lysosomal compartments. Representative images are shown from ≥2 experiments per condition with >3 worms per experiment.

In excess copper, the expansion compartment significantly increased in volume (Figure 1A,D). Compared to animals on control medium, animals on excess copper had an increase in total volume of the lysosome from ∼4.3 μm_3_ to ∼12.3 μm_3_ (∼2.9X increase) (Figure S1A, Table S1). The volume of the acidified compartment did not change significantly, whereas the volume of the expansion increased significantly from ∼0.73 μm_3_ to ∼10.1 μm_3_ (∼14X increase) (Figure S1C,E). In control animals, the expansion compartment represents ∼33% of the total volume of the lysosome - this increased significantly to ∼72% in copper excess (Figure S1B,D,F). These results suggest that excess copper may be stored in lysosomes.

In excess manganese, the expansion compartment significantly increased in volume (Figure 1A,E). Compared to animals on control medium, animals on excess manganese have an increased total volume of lysosomes from ∼4.3 μm_3_ to ∼13.2 μm_3_ (Figure S1A, Table S1). The volume of the acidified compartment did not change significantly, whereas the volume of the expansion increased significantly from ∼0.73 μm_3_ to ∼7.7 μm_3_ (∼10X increase) (Figure S1C,E). In control animals, the expansion compartment represents ∼33% of the total volume of the lysosome - this increased to ∼49% in manganese excess; this effect was not statistically significant with this sample size (Figure S1B,D,F).

Cadmium is not present in standard medium. Exposure to 200 μM cadmium appeared to cause lysosome remodeling suggestive of an increase in the volume of the expansion compartment (Figure 1A,F). The expansion compartments were typically shaped like a hemisphere. Compared to animals on control medium, animals on cadmium displayed an increase in total volume of the lysosome from ∼4.3 μm_3_ to ∼6.5 μm_3_ (∼1.5X increase); however, this trend was not statistically significant with this sample size (Figure S1A, Table S1). The volume of the acidified compartment did not change significantly, whereas the volume of the expansion increased from ∼0.73 μm_3_ to ∼4.2 μm_3_ (∼5.8X increase); however, this trend was not statistically significant with this sample size (Figure S1C,E). In control animals, the expansion compartment represents ∼33% of the total volume of the lysosome - this increased significantly to ∼62% in cadmium exposed animals (Figure S1B,D,F). These results suggest that exposure to cadmium triggers lysosome remodeling.

### Lysosomes contribute to detoxification of excess zinc, copper, and manganese

If lysosomes are a storage site for a metal, then we predict that reducing the number of lysosomes might have two effects: (1) Animals will become hypersensitive to metal excess, because lysosomes are not available for storage and detoxification. (2) Animals will become hypersensitive to metal deficiency, because lysosomes are not available as a source of stored metal. To test these predictions, we analyzed *glo-1(lf)* mutant animals that are characterized by a reduced number of intestinal lysosomes.^30^ We measured growth of larvae in 24 hours starting at the L4 stage, because the length of individual animals can be accurately and precisely quantified and serves as a good indicator of overall health. Wild-type and *glo-1(lf)* animals were cultured in a variety of concentrations of supplemental metal and metal chelators to test metal excess and deficiency, respectively. In the absence of supplements (0 μM), wild type and *glo-1(lf)* mutants displayed similar lengths, indicating that the *glo-1(lf)* mutants do not have a growth defect in standard conditions (Figure 2).

**Figure 2.**
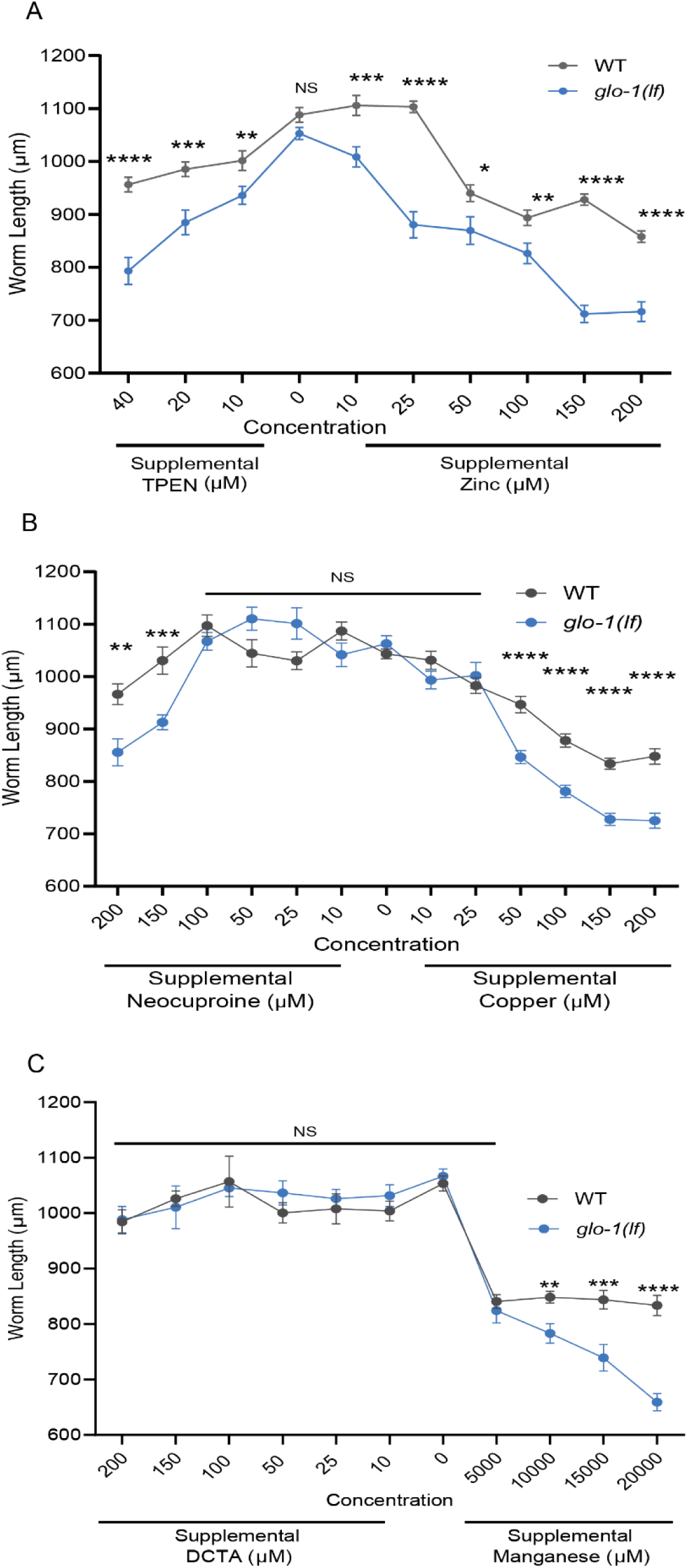
*glo-1(lf)* mutants with reduced numbers of lysosomes are hypersensitive to metal excess and deficiency. Wild type or *glo-1(zu391)* loss-of-function animals were synchronized at the L4 stage and cultured for 24 hours on standard NAMM medium or NAMM medium containing supplemental metal or chelator. The length of individual worms was measured using image analysis in FIJI. (A). NAMM medium containing the zinc chelator TPEN or supplemental zinc. Values represent the average length ± SEM. N=25-43 animals (wild type) and 30-46 animals (*glo-1(lf))*. (B) NAMM medium containing the copper chelator Neocuproine or supplemental copper. N=27-44 animals (wild type) and 26-44 animals (*glo-1(lf))*. (C) NAMM medium containing the manganese chelator DCTA or supplemental manganese. N=18-45 animals (wild type) and 23-47 animals (*glo-1(lf))*. Unpaired t-test: NS, not significant, * P ≤ 0.05, ** P ≤ 0.01, *** P ≤ 0.001, **** P ≤ 0.0001.

With low concentrations of supplemental zinc (10 and 25 μM), wild-type worms displayed normal growth, whereas *glo-1(lf)* mutants displayed significantly less growth (Figure 2A). In higher concentrations of supplemental zinc (50-200 μM), wild-type worms displayed reduced growth, whereas *glo-1(lf)* mutants displayed hypersensitivity (significantly less growth than wild type). Thus, *glo-1(lf)* animals are hypersensitive to both mild and extreme zinc excess. To create zinc deficient conditions, we utilized the zinc chelator N,N,N’,N’-tetrakis(2-pyridylmethyl)ethane-1,2-diamine (TPEN).^31^ Wild-type animals cultured on 10, 20, or 40 μM TPEN displayed reduced growth, indicating sensitivity to these levels of zinc deficiency, whereas *glo-1(lf)* mutants displayed hypersensitivity (significantly less growth than wild type) (Figure 2A). Thus, lysosomes promote resistance to both zinc excess and deficiency, consistent with the model that lysosomes are a storage site for zinc. These findings are consistent with previous results.^17,18^

With low concentrations of supplemental copper (10 and 25 μM), wild-type worms and *glo-1(lf)* mutants displayed similar growth (Figure 2B). In higher concentrations of supplemental copper (50-200 μM), wild-type worms displayed reduced growth, whereas *glo-1(lf)* mutants displayed hypersensitivity (significantly less growth than wild type) (Figure 2B). Thus, *glo-1(lf)* animals are hypersensitive to extreme copper excess. To create copper deficient conditions, we utilized the copper chelator Neocuproine.^32^ Wild-type animals and *glo-1(lf)* mutants cultured on 10-100 μM Neocuproine displayed relatively normal growth. When cultured with 150 or 200 μM Neocuproine, wild-type animals displayed reduced growth, indicating sensitivity to these levels of copper deficiency, whereas *glo-1(lf)* mutants displayed hypersensitivity (significantly less growth than wild type) (Figure 2B). Thus, lysosomes promote resistance to both copper excess and deficiency, consistent with the model that lysosomes are a storage site for copper.

With low concentrations of supplemental manganese (5,000 μM), wild-type worms and *glo-1(lf)* mutants displayed a similar reduction in growth (Figure 2C). In higher concentrations of supplemental manganese (10,000-20,000 μM), wild-type worms displayed reduced growth, whereas *glo-1(lf)* mutants displayed hypersensitivity (significantly less growth than wild type). Thus, *glo-1(lf)* animals are hypersensitive to extreme manganese excess. The results indicate that lysosomes promote resistance to manganese excess, consistent with the model that lysosomes are a storage site for manganese. To create manganese deficient conditions, we utilized the manganese chelator 1,2-Diaminocyclohexanetetraacetic acid monohydrate (DCTA).^33^ Wild type and *glo-1(lf)* mutant animals cultured on 10-200 μM DCTA did not display reduced growth, indicating both strains can tolerate this level of manganese deficiency (Figure 2C); these results do not test the model, because the highest levels of chelator did not cause a phenotype.

### Measurements of metals by X-ray Fluorescence Microscopy in isolated lysosomes

To investigate the hypothesis that lysosomes in intestinal cells store multiple metals, we developed a technique to quantify the metal content of isolated lysosomes. Previous studies in *C. elegans* have used the zinc dye FluoZin3 and the copper (I) dye CF4 to stain intact animals, which indicates that lysosomes accumulate zinc and copper, respectively.^16–18,29,34^ However, these dyes have limitations, and a manganese dye has not been employed. To quantitatively evaluate the metal content of lysosomes, we utilized the Bionanoprobe (BNP) at Argonne National Lab to conduct XFM.^35^ The BNP permits studies from biological samples at sub-100nm resolution, within a range of 4.5-35 KeV^36^ making it best suited to visualize metal content in lysosomes. XFM uses high energy X-rays to ionize inner electrons of atoms; in this unstable state, electrons from the outer shells drop down to fill those electron vacancies, thereby emitting energy in the form of X-ray fluorescence photons. Each element can be quantified in the same experiment based on its unique output. XFM has previously been used to analyze intact *C. elegans*^36,37^; to focus the analysis on lysosomes, we developed a method to dissect worms and expel intestinal lysosomes onto a grid suitable for XFM analysis. Transgenic animals were cultured in either control or excess zinc medium for 24 hours, and lysosomes were isolated by dissecting out the intestine, allowing the lysosomes to flow into the buffer, and pipetting the lysosomes onto a silicon nitride window for XFM analysis. To analyze the images, we measured the concentration of nine different elements inside the lysosome, subtracted background, and calculated the absolute number of each atom in the lysosome.

Each of the nine elements displayed a characteristic pattern of distribution within the spherical lysosome (Figure 3A-I, K-N). Phosphorus, calcium, and iron appeared at the perimeter of the lysosome, whereas sulfur, chlorine, potassium, manganese, copper, and zinc appeared to be distributed throughout the lumen of the lysosome. We calculated the average number of atoms in each lysosome; there were two general classes of abundance (Figure 3J). Phosphorus (1.6*10_9_ atoms per lysosome), sulfur (1.1*10_9_ atoms per lysosome), chlorine (6.0*10_8_ atoms per lysosome), potassium (1.5*10_9_ atoms per lysosome), and calcium (5.7*10_8_ atoms per lysosome) were relatively abundant, in the range of 10_9_ atoms per lysosome. By contrast, manganese (3.8*10_6_ atoms per lysosome), iron (4.5*10_6_ atoms per lysosome), copper (8.6*10_5_ atoms per lysosome), and zinc (7.8*10_6_ atoms per lysosome) were relatively less abundant, in the range of 10_7_ atoms per lysosome.

**Figure 3.**
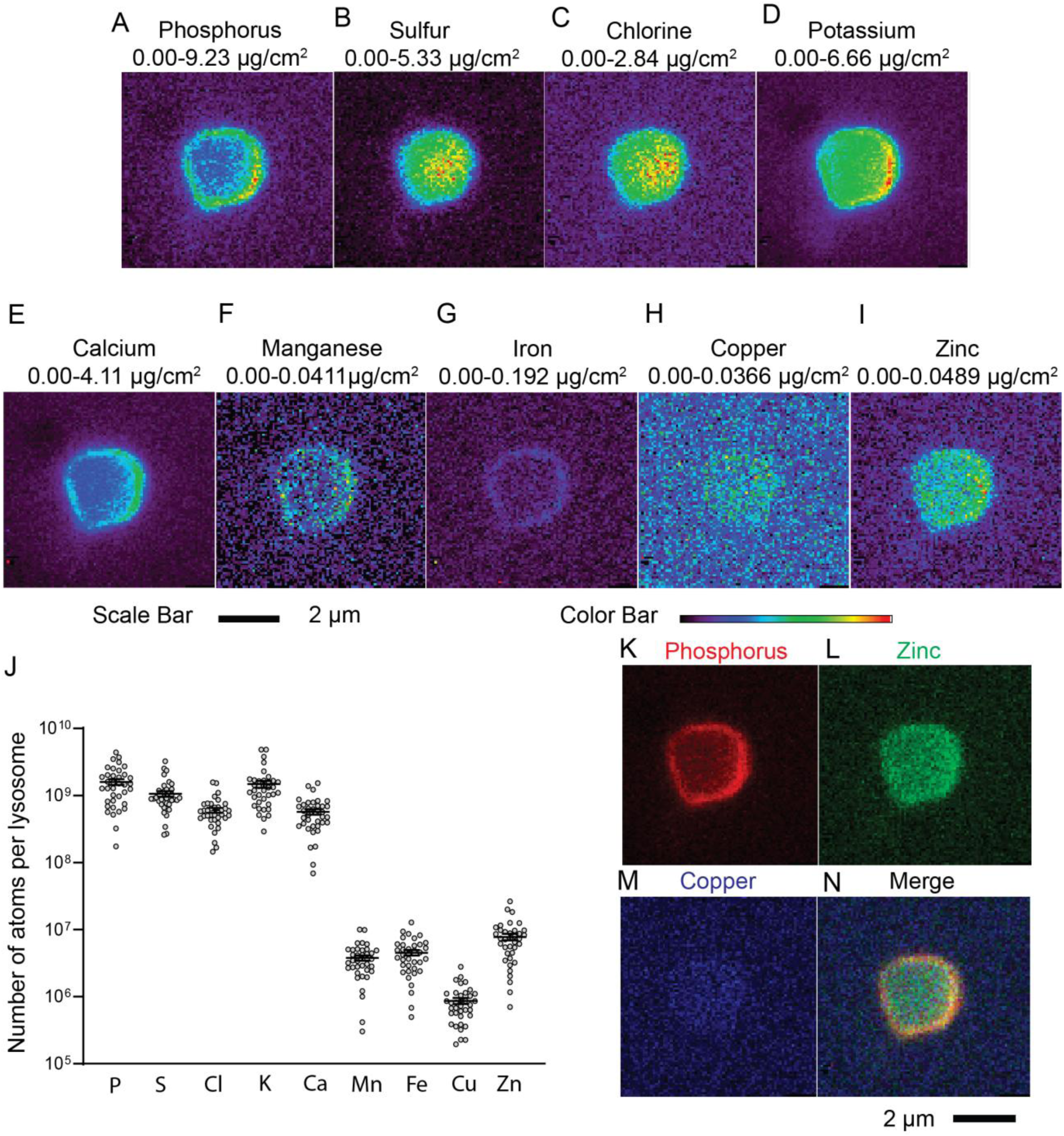
XFM analysis of individual lysosomes in standard medium. (A-I) Transgenic animals expressing CDF::GFP and ZIPT-2.3::mCherry were cultured on standard NAMM dishes. Intestinal lysosomes were isolated by dissection one day after the L4 stage, and XFM images were acquired with a 0.08μm step size and a dwell time of 500ms. Panels display XFM images from one representative lysosome, illustrating the distribution and intensity of phosphorus, sulfur, chlorine, potassium, calcium, manganese, iron, copper, and zinc. Color bar represents the spectrum of intensity; red is highest, and black is lowest. The extreme high and low intensity for each element is shown as the concentration range (μg/cm_2_). Scale bar = 2μm. (J) Each data point represents one lysosome; bars and whiskers represent the mean ± SEM. (N=37 lysosomes from 3 worms from one experiment). (K-N) False colored XFM images of phosphorous (red), zinc (green), and copper (blue); a merge displays overlaps in localization of these elements. The images are the same data as panels A, H, and I. Scale bar = 2μm.

To directly test the model that zinc is stored in lysosomes in response to dietary excess, we performed a similar analysis with 200 μM supplemental zinc in the medium (Figure 4). We quantified 60 individual lysosomes from multiple animals and compared the results to 37 animals cultured on control medium. Among the five abundant elements, the number of atoms per lysosome for phosphorus, sulfur, chlorine, and calcium was not significantly different from control, whereas potassium was significantly lower than control (1.8-fold reduction) (Figure S2A-E). Among the four less abundant transition metal elements, the number of atoms per lysosome for iron and copper was not significantly different from control, whereas manganese was significantly lower than control (2.6-fold reduction) (Figure S2F-H). For zinc, the number of atoms per lysosome increased significantly (13.8-fold increase), consistent with the model that zinc is stored in lysosomes in response to dietary excess (Figure 4K, Figure S2I).

**Figure 4.**
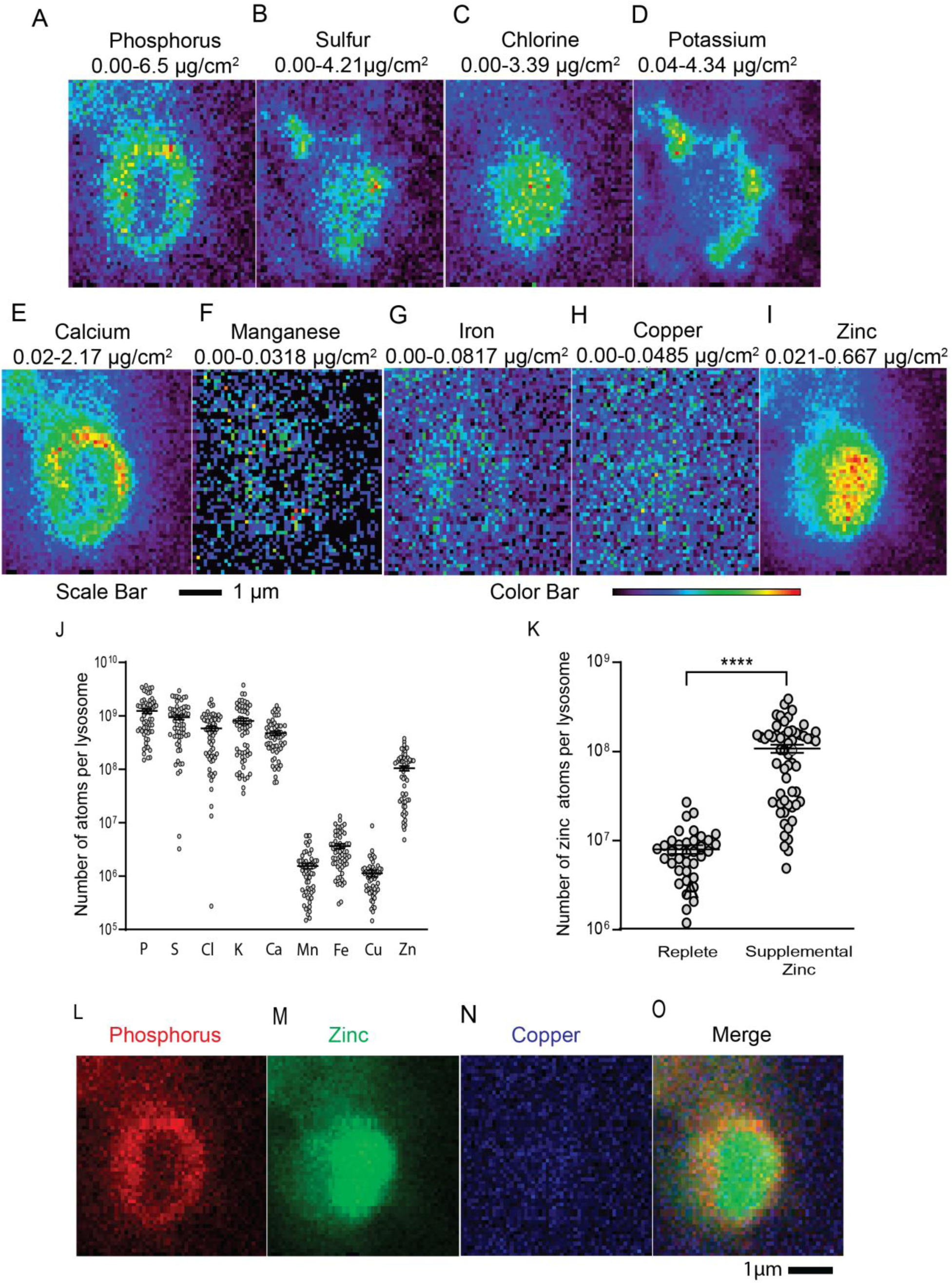
XFM analysis of individual lysosomes in excess zinc medium. (A-I) L4 stage transgenic worms expressing CDF::GFP and ZIPT-2.3::mCherry were transferred to NAMM dishes containing 200 μM supplemental zinc and cultured for 24 hours. Intestinal lysosomes were isolated by dissection from young adult animals, and XFM images were acquired with a 0.08μm step size and a dwell time of 500ms. Panels display XFM images from one representative lysosome, illustrating the distribution and intensity of phosphorus, sulfur, chlorine, potassium, calcium, manganese, iron, copper, and zinc. Color bar represents the spectrum of intensity; red is highest, and black is lowest. The extreme high and low intensity for each element is shown as the concentration range (μg/cm_2_). Scale bar = 1μm. (J) Each data point represents one lysosome; bars and whiskers represent the mean ± SEM. (N=60 lysosomes from 3 worms from one experiment). (K) Comparison of the number of zinc atoms per lysosome from animals cultured in standard NAMM (replete) or 200 μM supplemental zinc (excess) was performed by Welch’s t test. **** P ≤ 0.0001. (L-O) False colored XFM images of phosphorous (red), zinc (green), and copper (blue); a merge displays overlaps in localization of these elements. The images are the same data as panels A, H, and I. Scale bar = 1μm.

### The copper transporter CUA-1.1 localizes to both the acidified and expansion compartments

The *C. elegans cua-1.1* gene encodes a member of the ABC transporter family that functions as a copper transporter.^29^ CUA-1.1 protein localizes to the basolateral membrane of intestinal cells in replete copper, indicating it transports copper from the cytosol of intestinal cells into the pseudocoelomic space. However, in excess copper conditions, CUA-1.1 relocalizes to the membrane of lysosomes, indicating it transports copper from the cytosol of intestinal cells into the lumen of the lysosome to detoxify excess copper. These localization studies were performed using confocal microscopy, and the status of the expansion compartment was not reported.^29^ We hypothesized that excess copper would cause the expansion compartment to increase in volume as a result of delivering CUA-1.1 to the lysosome. To determine the localization of CUA-1.1 in the lysosome, we analyzed a transgenic strain that expresses CUA-1.1::GFP. In addition, we monitored copper (I) localization using the copper dye CF4.^38,39^ To validate the CF4 copper dye, we cultured wild-type animals with supplemental copper or the copper (I) chelator BCS. In copper (II) excess, lysosomes in intestinal cells displayed bright fluorescence with CF4; by contrast, in copper deficiency, the CF4 fluorescence was dim (Figure S3A-B). Quantification revealed a significant difference between these two conditions, indicating that the CF4 dye is copper (I) responsive in *C. elegans*, consistent with previous studies (Figure S3C).^29^

To resolve the acidified and expansion compartments, we performed super-resolution fluorescence microscopy. We analyzed two separate transgenic worm strains; one expresses CDF-2::GFP, which is known to localize to both the expansion compartment and the acidified compartment, whereas the other expresses CUA-1.1::GFP. Each strain was cultured on NAMM dishes with *E. coli* OP50 supplemented with LysoTracker blue and the copper dye CF4. In replete copper medium, lysosomes appear spherical with a very small expansion compartment that represents ∼13% of the volume (Figure S4, Table S2); CDF-2::GFP displayed an extensive overlap with LysoTracker and CF4 (Figure 5A). By contrast, CUA-1.1::GFP was primarily localized to the basolateral membrane and displayed weak staining of lysosomes; the spherical lysosomes displayed a very small expansion compartment that represents ∼7% on average of the volume. (Figure S4, Table S2); CUA-1.1::GFP displayed an overlap with LysoTracker and CF4 (Figure 5B). In excess copper, lysosomes displayed a significant increase in the volume of the expansion compartment to ∼66% of the total volume in the CDF-2::GFP strain and ∼74% of the total volume in the CUA-1.1::GFP strain (Figure S4). CDF-2::GFP localized to the acidified compartment and the expansion compartment. By contrast, LysoTracker and CF4 localized to the acidified compartment but not the expansion compartment (Figure 5D,E). In excess copper, CUA-1.1::GFP displayed a similar pattern to CDF-2::GFP; CUA-1.1::GFP localized to acidified compartment and the expansion compartment, whereas CF4 and LysoTracker localized to the acidified compartment but not the expansion compartment (Figure 5D,F).

**Figure 5.**
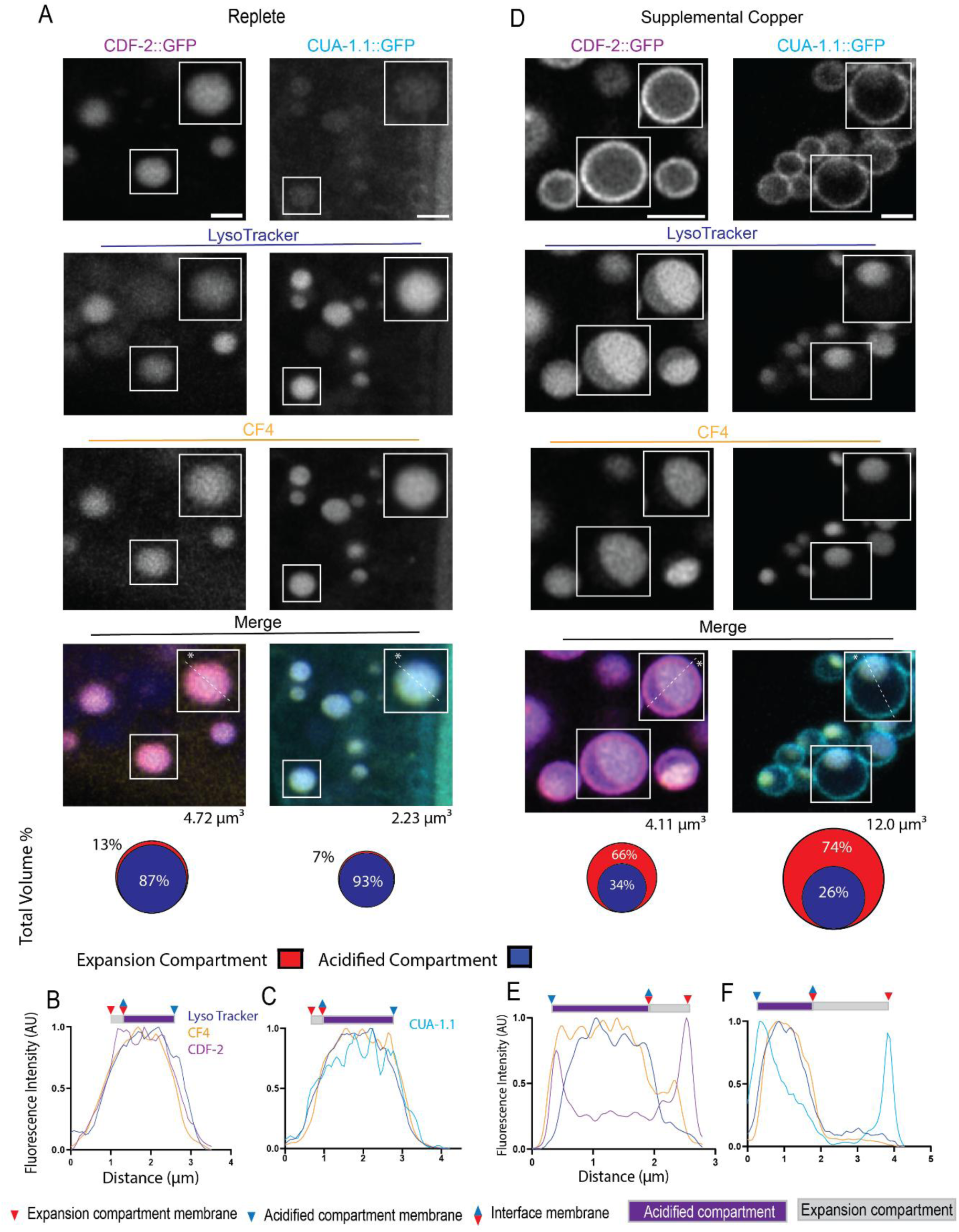
The copper transporter CUA-1.1 localizes to the expansion compartment and acidified compartment membranes in excess copper. (A) Transgenic L4 stage animals expressing CDF-2::GFP (magenta) or CUA-1.1::GFP (cyan) were cultured for 24 h in LysoTracker Blue (blue) and the copper dye CF4 (yellow) on either NAMM medium (replete), or NAMM medium containing 100μM supplemental copper, and imaged by super-resolution microscopy. Images show a portion of the intestine of 1-day old adult animals. White boxes are enlarged in the upper right of each image. Scale bar = 2 µm. Dashed white lines and asterisks indicate the lysosome analyzed below. Cartoons illustrate the average total volume of a lysosome (μm_3_) and the percent represented by the acidified (blue) and expansion (red) compartments (Figure S4, Table S2). (B-F) Line scans of individual lysosomes in replete (B,C) or excess copper (E,F). Fluorescence intensity along the path of the line was measured beginning at the white asterisk. CDF::GFP, CUA-1.1::GFP, LysoTracker, and CF4 are displayed as magenta, cyan, blue, and yellow respectively. For each color line, the highest value of fluorescence intensity is set to 1.0 in arbitrary units (AU), and the lowest fluorescence intensity is set to 0. All other values are normalized. Triangles indicate positions of membranes, and rectangles indicate lysosomal compartments. Representative images of N=3 worms per condition.

### CUA-1.1 and CDF-2 localize to the same lysosomes in intestinal cells

*C. elegans* intestinal cells contain many lysosomes. One possible model is that zinc storing lysosomes are separate from copper storing lysosomes; alternatively, the same lysosomes may store both zinc and copper. To distinguish these models, we generated a transgenic strain that expresses both CUA-1.1::GFP and CDF-2::mCherry. In copper replete medium, super-resolution microscopy revealed that CUA-1.1::GFP localized primarily to the basolateral membrane and weakly to the lysosomal membrane; by contrast, CDF-2::mCherry localized specifically to lysosomes (Figure 6A). The same lysosomes that were strongly fluorescent for CDF-2::mCherry were also weakly fluorescent for CUA-1.1::GFP, demonstrating that both transporters reside on the same lysosomes in replete medium (Figure 6B, C). To determine the effect of excess copper, we cultured these animals in supplemental copper for 24 hours with *E. coli* OP50 containing LysoTracker. Super-resolution microscopy revealed that lysosomes were remodeled; the percent of the lysosome volume occupied by the expansion compartment increased from ∼30% in copper replete medium to ∼67% in excess copper medium (Figure S5, Table S3). Importantly, CDF-2::mCherry and CUA-1.1::GFP co-localized to both the acidified and expansion compartments of the same lysosomes (Figure 6B, D). In these conditions, the two transporters displayed a highly overlapping pattern. These results indicate that the same lysosomes contain zinc and copper transporters and likely store both metals.

**Figure 6.**
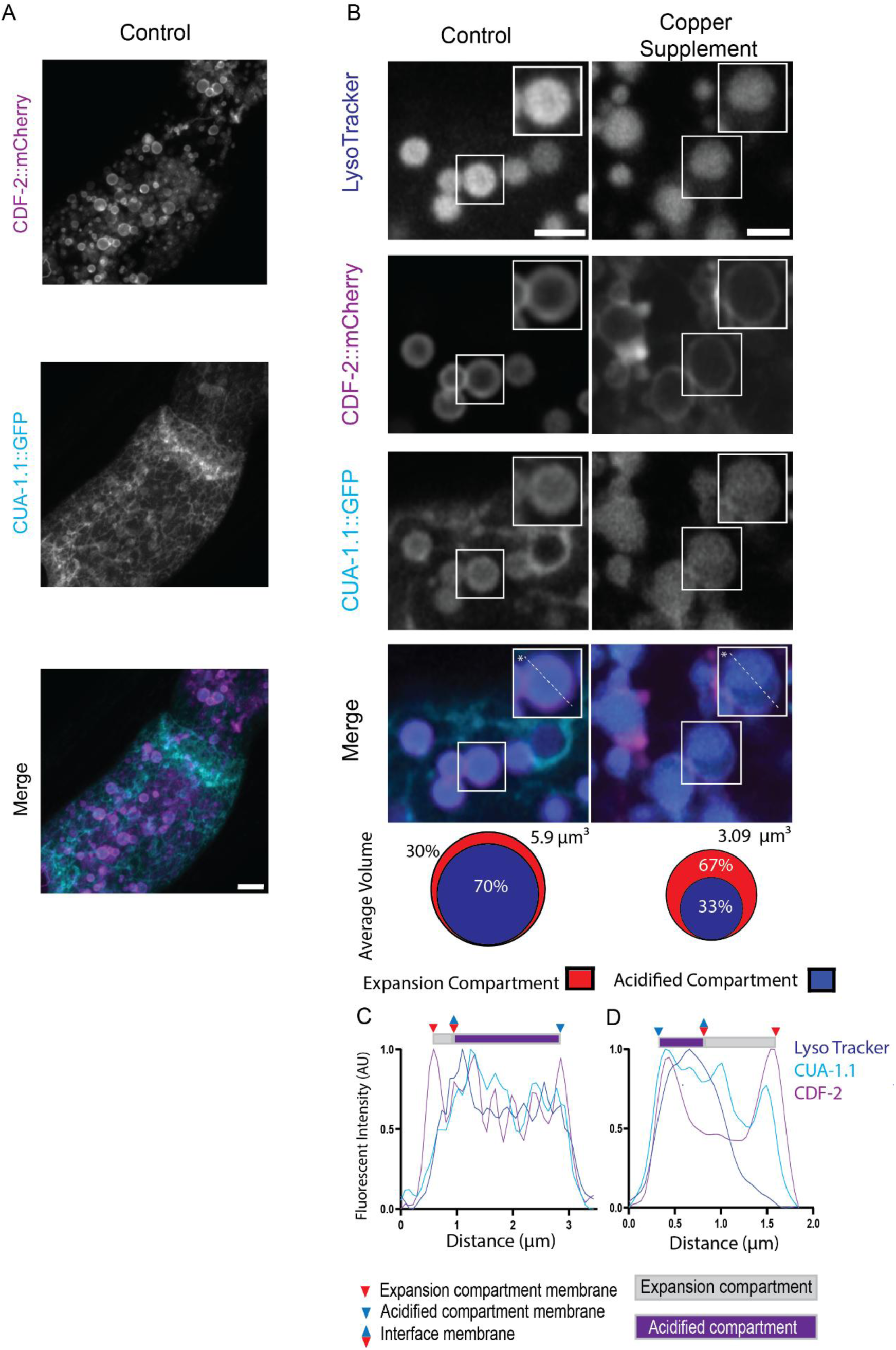
CDF-2 and CUA-1.1 localize to the same lysosomes. (A) Transgenic L4 stage animals expressing CDF-2::mCherry (magenta) or CUA-1.1::GFP (cyan) were cultured for 24 h on NAMM medium (control) and imaged by super-resolution microscopy. Images show a portion of the intestine of 1-day old young adult animals. Scale bar = 2 µm. (B) The same strain was cultured for 24 h in LysoTracker Blue (blue) on either NAMM medium (control), or NAMM medium containing 100μM supplemental copper (copper supplement), and imaged by super-resolution microscopy. Images show a portion of the intestine of 1-day old young adult animals. White boxes are enlarged in the upper right of each image; Scale bar = 2 µm. Dashed white lines and asterisks indicate the lysosome analyzed below. Cartoons illustrate the average total volume of a lysosome (μm_3_) and the percent represented by the acidified (blue) and expansion (red) compartments (Figure S5, Table S3). (C-D) Line scans of individual lysosomes in replete (C) or excess copper (D). Fluorescence intensity along the path of the line was measured beginning at the white asterisk. CDF::mCherry, CUA-1.1::GFP, and LysoTracker are displayed as magenta, cyan, and blue, respectively. For each color line, the highest value of fluorescence intensity is set to 1.0 in arbitrary units (AU), and the lowest fluorescence intensity is set to 0. All other values are normalized. Triangles indicate positions of membranes, and rectangles indicate lysosomal compartments. Representative images of N=3 worms per condition.

## Discussion

Transition metals are essential for all organisms because of their versatile chemical properties. However, this addiction to metals has a downside – excess metals are toxic and must be managed carefully. In the case of iron, excess metal is stored in the cytoplasm in a complex with ferritin^40,41^. Lysosomes are emerging as an important site of excess metal detoxification and storage; these lysosomes also provide a source of metal in response to deficiency. Here we used *C. elegans* intestinal cells to probe the generality of excess metal storage in lysosomes by analyzing zinc, copper, manganese, and cadmium. In *C. elegans,* analysis with the zinc dye FluoZin-3 indicated that intestinal lysosomes store excess zinc.^17^ Super-resolution microscopy revealed that lysosomes contain two compartments, a LysoTracker positive acidified compartment, and a LysoTracker negative expansion compartment.^18^ The zinc transporters CDF-2 and ZIPT-2.3 localize to the acidified compartment, whereas CDF-2 but not ZIPT-2.3 localizes to the expansion compartment. In excess zinc, CDF-2 mRNA and protein are upregulated, and CDF-2 positive vesicles fuse to the expansion compartment, resulting in an increase in volume and the amount of CDF-2 protein. Based on these observations, Mendoza et al (2024)^18^ proposed that the increase in the volume of the expansion compartment is caused by the fusion of vesicles delivering CDF-2 protein. To test the hypothesis that the increase in the expansion compartment volume is a general mechanism to deliver metal transporters and increase the capacity to store metals, we investigated additional metals. In addition, we developed a technique to quantify metals in isolated lysosomes using X-ray Fluorescence Microscopy. In replete zinc medium, we measured 7.8*10_6_ zinc atoms per lysosome, which appears to be distributed throughout the lumen of the lysosomes. In excess zinc medium, the amount increased dramatically to 1.0*10_8_ zinc atoms per lysosome. This result is consistent with the conclusion based on the dye FluoZin-3 and formally demonstrates that lysosomes are an important site of zinc storage during excess.

The observation that labile zinc (visualized by FluoZin-3) was absent in the expansion compartment, despite CDF-2 localization suggested that the zinc dye was unable to access bound zinc in the expansion compartment.^18^ XFM revealed that total zinc was detected throughout the entire lysosome, supporting the idea that CDF-2 could be importing zinc into the expansion compartment as well as the acidified compartment. Developing an XFM procedure to isolate lysosomes is a novel approach and is necessary to understand how lysosomes acquire zinc and uncover the elements that are present within them. Previous XFM analyses in worms were performed on whole adults and isolated embryos,^37,42^ however here we demonstrate the ability to isolate individual lysosomes from intestines. This novel innovation opens opportunities for refining metal detection in organelles and correlating zinc content with compartments.

The heavy metal cadmium accumulates in animals.^43,44^ However, because cadmium does not have a physiological function in animals, this accumulation is not an evolved response to store excess metal for future use. Earley et al (2021)^23^ reported that cadmium hijacks the high zinc response by binding and activating HIZR-1, which results in the upregulation of CDF-2. Here we show that 200μM cadmium causes lysosomal remodeling. We can interpret this response as an upregulation of genes that contributes to the growth of the expansion compartment. Cadmium specific dyes have not been reported, so it is unclear where cadmium accumulates in *C. elegans*.

The physiological metal manganese has been studied in *C. elegans*.^21,44^ Because manganese specific dyes are not readily available, the localization of manganese in *C. elegans* is not well established. Here we show that excess manganese causes lysosome remodeling characterized by an increase in the volume of the expansion compartment. We speculate that this reflects the delivery of vesicles containing a manganese transporter to the expansion compartment of the lysosome to increase the capacity for manganese storage. However, this manganese transporter has not been identified. Consistent with this model, XFM revealed ∼1.5*10_6_ and 3.5*10_6_ manganese atoms per lysosome in replete and excess zinc medium, respectively. To evaluate the physiological importance of lysosomes for excess manganese detoxification, we analyzed *glo-1* mutants that have a reduced number of lysosomes. These mutants were hypersensitive to excess manganese, suggesting that lysosomes sequester manganese. Together, these results indicate that lysosomes in intestinal cells of *C. elegans* play an important role in manganese storage. Important future experiments to test this hypothesis include the identification of the transporters that store manganese in lysosomes during excess and release manganese from lysosomes during deficiency.

Copper is an essential metal; Chun et al (2017)^29^ proposed that copper is stored in lysosomes in intestinal cells based on the localization of the copper dye CF4 to lysosomes and the localization of the copper transporter CUA-1.1 to lysosomes in copper excess. Here we confirmed these findings with the copper dye CF4, and we extended this analysis by performing XFM on isolated lysosomes. We detected 8.6*10_5_ and 1.1*10_6_ copper atoms per lysosome in replete and excess zinc medium, respectively. In addition, excess copper caused lysosomal remodeling characterized by an increase in the volume of the expansion compartment. This result suggests that a copper transporter is delivered to the expansion compartment by vesicle fusion. To evaluate the physiological significance of copper storage in lysosomes, we analyzed *glo-1* mutants with a reduced number of lysosomes. These mutants were hypersensitive to both copper excess and deficiency, suggesting that copper storage in lysosomes detoxifies excess copper, and copper is mobilized from lysosomes during copper deficiency. These results raise an important conceptual question – do the same lysosomes store both copper and zinc, or are these metals stored in separate lysosomes? To address this question, we performed colocalization studies of the zinc transporter CDF-2 and the copper transporter CUA-1.1. In copper replete medium, CUA-1.1 is localized primarily to the baso-lateral membrane of intestinal cells, whereas CDF-2 is constitutively localized to the membrane of lysosomes. Thus, in these conditions the two transporters display minimal colocalization. However, in copper excess conditions, CUA-1.1 and CDF-2 colocalize to the same lysosomes, which appear to be essentially all the LysoTracker positive lysosomes in the intestinal cells. Furthermore, CUA-1.1 and CDF-2 colocalize to both the expansion compartment and the acidified compartment. These results indicate that all lysosomes in intestinal cells store both copper and zinc. We speculate that the expansion compartment increases in volume in excess copper due to the fusion of vesicles that contain CUA-1.1. These vesicles likely originate from the baso-lateral plasma membrane, where CUA-1.1 is localized in copper replete conditions. Roh *et. al. (*2012)^17^ reported that PGP-2 protein localizes to the expansion compartment, so CUA-1.1 is the third protein reported to localize to the expansion compartment. Indeed, earlier work combining fluorescent copper probes and XFM reveal copper fluxes in neurons consistent with vesicle transport.^45^ Recent studies are also beginning to demonstrate the emerging role of lysosomes in copper homeostasis. For example, mammalian studies have recently shown that lysosomal solute carrier 46a3(SLC56A3) mediates copper homeostasis in the liver.^46^ Furthermore, copper homeostasis also seems to be influential in the response against pathogens by utilizing copper transporter ATP7A to take down phagasomes.^47^

Exposure to a battery of transition metals suggests that there is morphological diversity in remodeling. Zinc exposure leads to lysosomes that display an acidified compartment that projects outward and an expansion compartment that is extremely extended (Figure 1). Exposure to manganese and copper revealed morphological changes in which an acidified compartment appeared encapsulated within the expansion compartment. Lysosomes exposed to cadmium showed an increase in the expansion compartment and a decrease of the acidified compartment. These observations indicate that lysosomes may be an important site for stress response but also a potential storage place for toxic transition metals. Transition metal exposure seems to yield different spatial relationships between compartments. These results lead us to speculate that other metal ion transporters contribute to these morphological changes and ask what their counter ions could be. Furthermore, it is unclear if these morphological changes are representative of a “stable” state of a lysosome or if the morphological shifts represent a “transient” stage.

Our discovery of metal-responsive expansion compartments raises questions about their evolutionary conservation. Recently, Tavakoli *et al*. (2025)^48^ identified similar structures termed “hemifusomes” in human cells, and they propose that hemifusomes are an intermediate step toward the development of multivesicular bodies. While lysosomes are already known to undergo morphological remodeling including tubulation and spherical remodeling,^49,50^ our work shows that a distinct expansion compartment has a function in transition metal homeostasis and represents a novel biological role for lysosomes that leverages a compartment. Based on our findings, we developed a model that describes copper and zinc management by lysosomes (Figure 7). The expansion compartment is always present, but it is contracted and difficult to visualize in standard media. In copper replete medium, CUA-1.1 is localized to the basolateral membrane, whereas CDF-2 is localized to the lysosomes. As the organism transitions to copper excess, CUA-1.1 containing vesicles depart the basolateral membrane and fuse with the expansion compartment, causing it to increase in volume and increase the capacity for copper storage in the lysosomes. Increased copper storage in lysosomes reduces the cytosolic concentration of copper, thereby reducing toxicity. CDF-2 and CUA-1.1 reside on the same lysosomes, which store both copper and zinc. Our results raise the possibility that many metals, and perhaps some organic molecules, can be stored and released from the same lysosomes through the parallel action of specific transporters.

**Figure 7.**
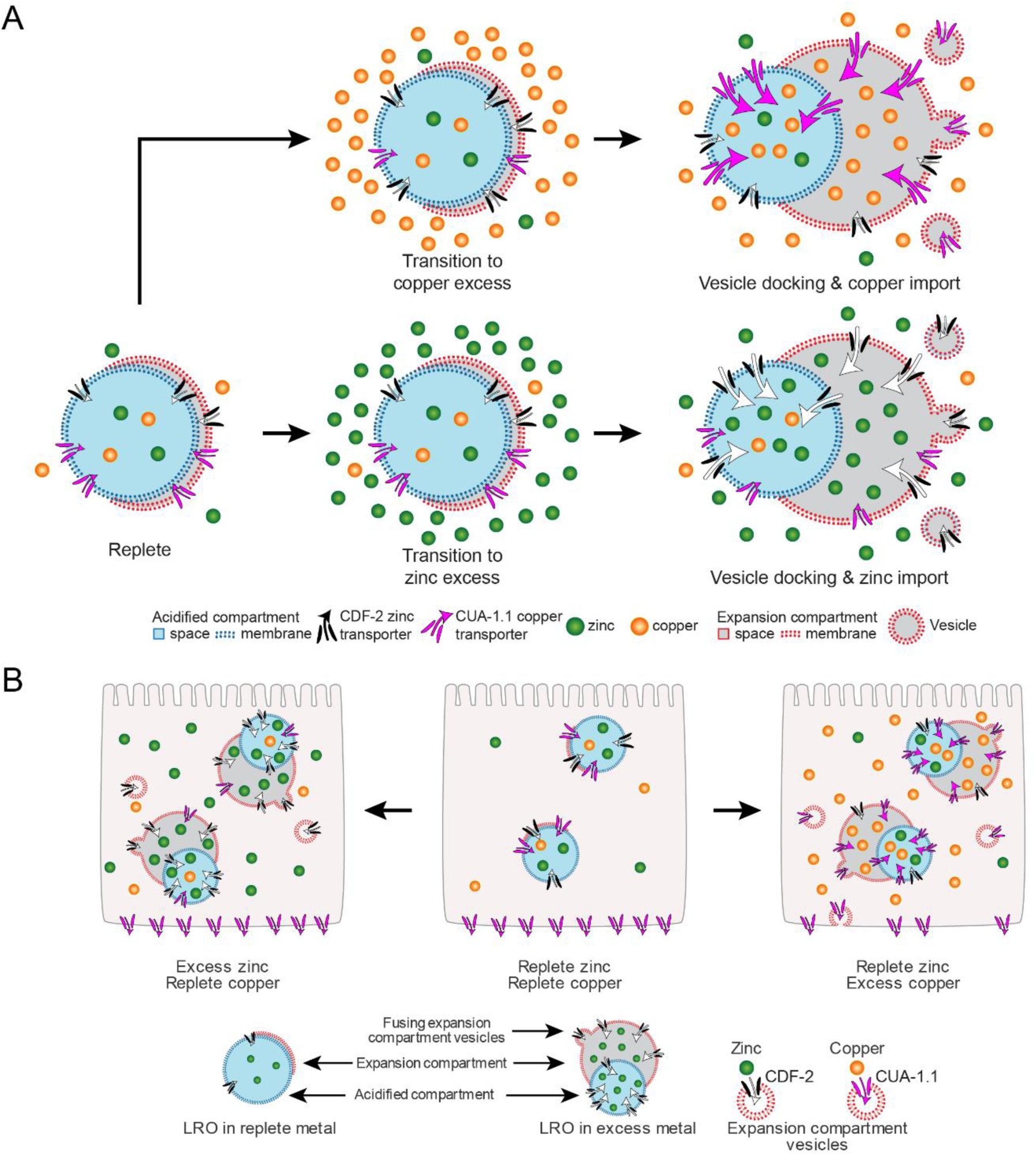
A model of lysosome remodeling during excess zinc and copper. (A) In replete medium, the lysosome has a high-volume acidified compartment (blue), a low-volume expansion compartment (red), and a small number of zinc and copper transporters. In excess copper (orange balls), vesicles containing the copper transporter CUA-1.1 (magenta) fuse to the expansion compartment, increasing the volume, number of copper transporters, and the capacity for copper storage. In excess zinc (green balls), vesicles containing the zinc transporter CDF-2 (white) fuse to the expansion compartment, increasing the volume, number of zinc transporters, and the capacity for zinc storage. While both cases lead to an increase in the volume of the expansion compartment, the transporter delivered by the expansion compartment vesicles is different. (B) In replete medium (center), an intestinal cell (pink) contains lysosomes with a low-volume expansion compartment. The copper transporter CUA-1.1 is primarily localized to the basolateral membrane, whereas the zinc transporter CDF-2 is primarily localized to the lysosome membrane. In excess zinc (left), expression of *cdf-2* mRNA and protein increases, and expansion compartment vesicles containing CDF-2 fuse to the expansion compartment to increase the volume and capacity for zinc storage. In excess copper (right), expansion compartment vesicles transfer CUA-1.1 from the basolateral plasma membrane to the lysosome membrane, increasing the volume of the expansion compartment and the capacity for copper storage.

## Experimental Procedures

### C. elegans Strains

*C. elegans* strains were cultured on nematode growth medium (NGM) dishes at 20ᵒC with OP50 *E. coli* unless otherwise stated. To manipulate metal concentrations, we used Noble Agar Minimal Media (NAMM) because it limits precipitation of metals.^51^ The wild-type strain and parent of all mutant strains was Bristol N2.^52^ Loss-of-function mutations were *cdf-2(tm788)*^16^ *and glo-1(zu391)* X.^30^ Extrachromosomal or integrated arrays were used to tag CDF-2, ZIPT-2.3, and CUA-1.1: *amEx342* is a multicopy extrachromosomal array that expresses CDF-2 fused to mCherry (*cdf-2::mCherry;rol-6_D_*).^17^ *amIs4* is an integrated multicopy array that expresses CDF-2 fused to GFP (*cdf-2(tm788);amIs4).*^16^ Extrachromosomal array *amEx193*, contains *zipt-2.3* fused to mCherry.^18^ The following strains were used: WU1984 contains *cdf-2(tm788); amIs4; amEx193;Pmyo2::mCherry::unc-54utr*^18^, BK014 contains P*_vha-6_* ::CUA-1.1::GFP::unc-54 3’ UTR. WU2132 contains *cdf-2 amEx342rol^D^*; P*_vha-6_* ::CUA-1.1::GFP::unc-54 3’ UTR. WU2132 was created by crossing parental *cdf-2 amEx342rol_D_* hermaphrodites with male P*_vha-6_* ::CUA-1.1::GFP::unc-54 3’ UTR worms. Worms containing both *cdf-2 amEx342rol^D^* and P*_vha-6_* ::CUA-1.1::GFP::unc-54 3’ UTR were tracked across three generations.

### Transition Metal and Chelator Supplementation

Metals and metal chelators were all dissovled in MilliQ H_2_O or ethanol. Stock solutions of metals were added to molten NAMM media to yield a final concentration of 10-200μM supplemental zinc (ZnSO_4,_ Sigma Aldrich), 10-200μM supplemental copper (CuCl_2_, Sigma Aldrich), 5-20mM supplemental manganese (MnCl_2_, Sigma Aldrich), or 200μM cadmium (CdCl_2_ Sigma Aldrich). Stock solutions of chelators were added to molten NAMM media to yield a final concentration of 10-200 μM of the copper chelator Neocuproine (Sigma Aldrich),10-40 μM of the zinc chelator N,N,N’,N’-tetrakis(2-pyridylmethyl)ethane-1,2-diamine (TPEN, Sigma Aldrich) or 10-200 μM of the manganese chelator 1,2-Diaminocyclohexanetetraacetic acid monohydrate (DCTA, Sigma Aldrich).

### Worm Growth Experiments

Synchronized wild type and *glo-1(zu391)* L4 stage worms were transferred to control Noble Agar Minimum Media (NAMM) dishes or NAMM dishes that were supplemented with the copper chelator Neocuproine, the zinc chelator TPEN, the manganese chelator DCTA, CuCl_2_, ZnSO_4_, or MnCl_2_. Worms were cultured for 24 hours on dishes seeded with *E. coli* OP50. Dishes were placed on a Leica stereomicroscope, and images of individual worms were captured. Images were analyzed for individual worm lengths using FIJI (NIH) software by measuring from the tip of the nose to the tip of the tail for each worm.

### Super-Resolution Microscopy

L4 stage animals expressing CDF-2::GFP and ZIPT-2.3::mCherry, or CDF-2::mCherry and CUA-1.1::GFP, were cultured for 24 hours on NAMM dishes containing 200μM zinc, 100μM copper, 20mM manganese, or 200μM cadmium. To visualize copper, we diluted the copper dye CF4 into *E. coli* OP50 to obtain a final concentration of 25 μM.^38^ To visualize lysosomes, we diluted LysoTracker Blue DND-22 dye (Thermofisher) in *E.coli* OP50 to a final concentration of 1 μM. Worms consumed these dyes with their bacterial food. After 24 hours, the worms were picked into 50 mM NaN_3_ in MilliQ that was pipetted onto an agar pad. The agar pad was sealed with Vaseline to a coverslip. Super-resolution microscopy was performed with the Zeiss LSM 880 II Confocal with Airyscan. Individual lysosomes were selected for analysis if they displayed all relevant signals being tested in each experiment, and they were isolated from neighboring lysosomes. Lysosome images were captured in z-stack using the 60x objective. CDF-2::GFP and CUA-1.1::GFP were detected using the 488 nm laser, LysoTracker Blue DNDdye was detected using the 405 nm laser, and ZIPT-2.3::mCherry, CDF-2::mCherry and CF4 dye were detected using the 561 nm laser. Images were deconvolved using AiryScan processing to achieve ∼120 nm resolution.

### Image Analysis

#### Line Scans

Line scans were performed on images of lysosomes from transgenic worms expressing CDF::GFP and ZIPT-2.3::mCherry, CDF-2::GFP, CUA-1.1::GFP, and CDF-2::mCherry and CUA-1.1::GFP. Lysosomes were selected for analysis if the z-stack encompassed the entire lysosome, and if they displayed GFP, mCherry, and LysoTracker (Figure 1,6) or GFP, LysoTracker, and CF4 (Figure 5). In FIJI, lines were drawn across a single center slice of a whole lysosome starting from the acidified compartment and proceeding through the expansion compartment. For control lysosomes that have a small expansion compartment, the orientation was chosen at random. The intensity of GFP, mCherry, CF4, and LysoTracker was plotted as fluorescence intensity on the y axis. The line drawn across the lysosome forms the x axis, displaying distance. For each channel, the maximum fluorescence intensity is set to 1.0 in arbitrary units (AU), and the lowest fluorescence intensity is set to 0. All other values are normalized.

Measurements of fluorescence intensity for CF4 (Figure S3) were done in FIJI. An ROI was drawn around individual lysosomes, and the average fluorescence intensity of the ROI was reported in arbitrary units (AU).

#### Volume Calculation

The diameter (d) of each expansion and acidified compartment was measured in FIJI by drawing a line across each compartment three times and calculating the average value. In control, supplemental zinc, supplemental copper, and supplemental manganese, compartments were assumed to be spherical. Based on this assumption and the measured diameters, we calculated the radius as *r*= *d*/2 for each compartment and then calculated the volume as V= 4/3 *πr*^3^. In control, supplemental zinc, supplemental copper, and supplemental manganese, the acidified compartment appears to be contained within the expansion compartment (a sphere within a sphere). To calculate the volume of the expansion compartment alone, we subtracted the volume of the acidified compartment from the volume of the total lysosome (V_E_=V_T_-V_A_). For the cadmium exposure experiments, the expansion compartment appears to have the shape of a cap on a sphere instead of a sphere itself. Therefore, we calculated the total volume of the lysosome by measuring the diameter three times and then calculating the average value. The total volume of the sphere was calculated as V= 4/3 *πr*^3^. To determine the volume of the expansion compartment, we measured the volume of the cap on the sphere with the equation 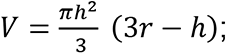 we measured r three times to determine the average. To determine the volume of the acidified compartment, we subtracted the expansion compartment volume from the total volume (V_A_=V_T_-V_E_).

### X-Ray Fluorescence Microscopy Sample preparation, Scanning parameters, and Data Analysis

#### Preparation of isolated lysosomes

L4 stage worms were picked from dishes with mixed populations and cultured on control and zinc supplemented NAMM dishes for 24 hours, during which they develop into young adults. A microscope slide was set up to accommodate three drops of Milli-Q water and one drop of 100 mM ammonium acetate buffer. None of the drops were allowed to touch each other to avoid contamination. Worms were successively transferred by picking into each water drop to wash off bacteria and then transferred by picking into the drop of ammonium acetate buffer. Using a needle, worms were cut to extrude the intestine. Lysosomes were allowed to flow into the buffer. The isolated lysosomes were pipetted and transferred to a Silicon Nitride window (Silson) using a 150 μm oocyte striper tip (Cooper Surgical) and allowed to air dry.

#### XFM

Isolated lysosomes were raster scanned with a pixel size of 80 nm and a dwell time of 500ms per pixel. A double-crystal Si <111> monochromator and a pair of Fresnel zone plates were used to deliver a 10 keV focused beam of ∼80 nm. All XFM experiments were performed at the Bionanoprobe^53^ at the Advanced Photon Source (Argonne National Laboratory, Argonne IL).

#### Data Analysis

All data were analyzed using the MAPS software.^54^ ROIs of individual lysosomes were traced in MAPS, and quantities obtained from whole individual lysosomes were generated in MAPS in femtograms. To calculate the total elemental content for each ROI, we subtracted the background from the initial femtograms to generate the background subtracted mass. The mass was then converted to atoms/lysosome.

#### Statistical analysis

For all XFM data comparisons were performed using the two-tailed unpaired Student’s t-test, or a 2-way ANOVA, and a P value <0.05 was considered statistically significant (Figure S2).

## Statistical analysis

Statistical tests were either a One-Way Anova (Figure S1), Welches t-test (Figure 4, Figure S3, S4, and S5) or the two-tailed unpaired t-test (Figure 2). P value <0.05 was considered statistically significant.

## Acknowledgements

We thank Daniel Herrera for initiating copper experiments. pCFJ90-Pmyo-2::mCherry::unc54utr was a gift from Erik Jorgensen (Add gene plasmid #19327). NIH R01GM068598 to K.K., 1K99GM146016-01 and 4R00GM146016 to A.D.M. R01GM79465 to C.J.C. C.J.C. is a CIFAR Fellow. J.R.A. was supported by T32GM142607. Confocal/super-resolution microscopes were purchased with support from the Washington University School of Medicine Office of Research Infrastructure Programs, part of the NIH Office of the Director (grant OD021629). The synchrotron X-ray measurement was performed the Advanced Photon Source, a U.S. Department of Energy (DOE) Office of Science user facility at Argonne National Laboratory and is based on research supported by the U.S. DOE Office of Science-Basic Energy Sciences, under Contract No. DE-AC02-06CH11357. The Bionanoprobe development was supported by NIH ARRA grant SP0007167.

## Author contributions

A.D.M. and K.K. designed the research; A.D.M., S.T., R.H., S.H., and D.L.S., performed research; B.-E.K., A.T.P and C.J.C. contributed new reagents; S.C and S.V. analytic tools; A.D.M., K.R., M.J., M.T.G., C.R., and J.R.A., analyzed data; J.R.A. and H.L., generated figures, and A.D.M., J.R.A., and K.K. wrote the manuscript.

---

**Supplemental Figure 1 (with main Figure 1).**
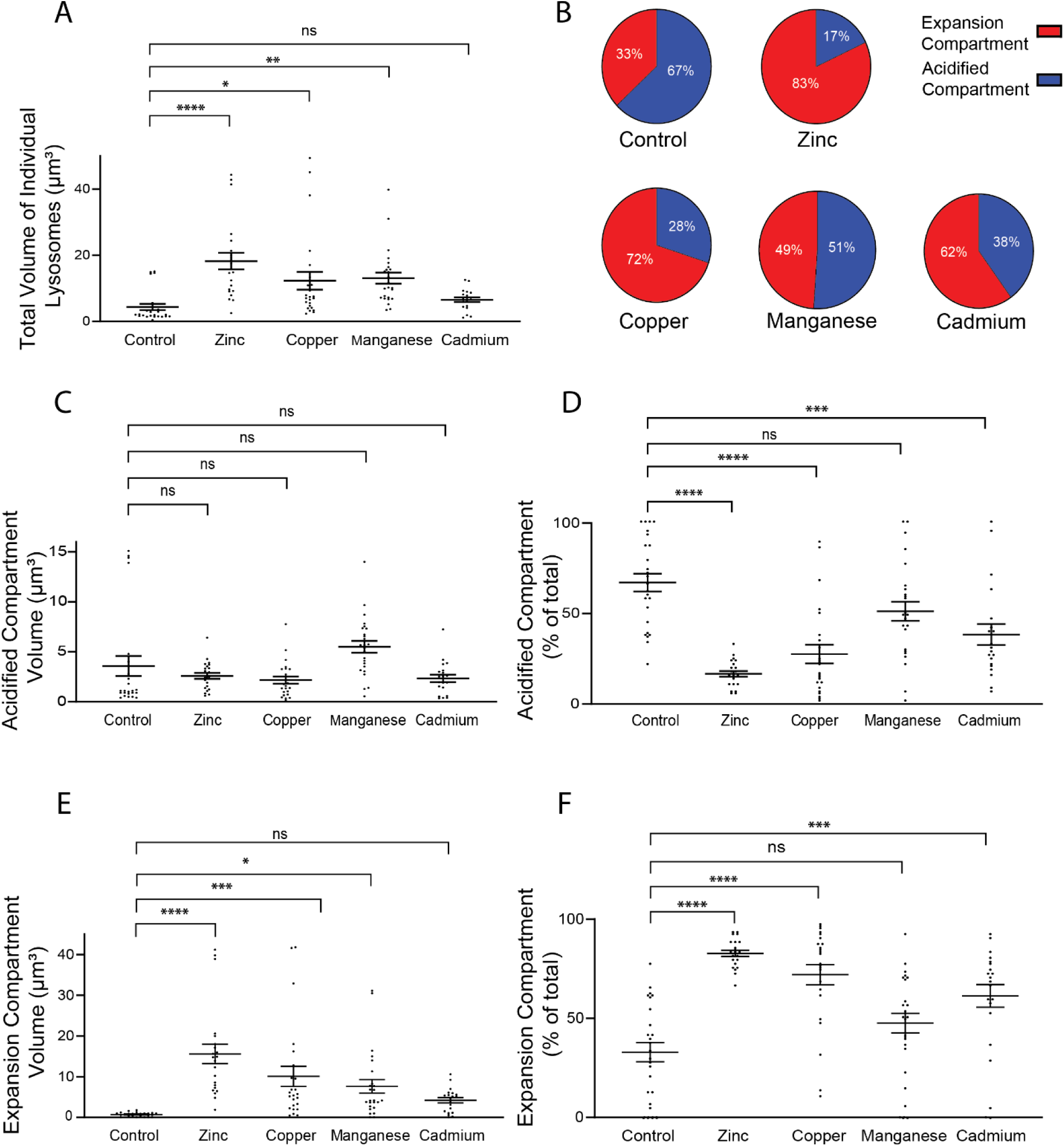
Quantification of acidified compartment volume, expansion compartment volume, and total lysosome volume. L4 stage transgenic animals expressing CDF-2::GFP and ZIPT2.3::mCherry were cultured for 24 hours in LysoTracker blue in NAMM medium (Control), or NAMM medium containing 200 μM supplemental zinc, 100μM supplemental copper, 20mM supplemental manganese, or 200μM cadmium. Lysosomes were imaged at the young adult stage by super-resolution microscopy with green, red, and blue fluorescence. The volumes of the expansion compartment and the acidified compartment were calculated from 3-dimensional images by assuming the compartments are shaped like spheres or hemispheres (see Experimental Procedures for details). (A, C, E) Total lysosomal, acidified compartment, and expansion compartment volume (μm_3_). (B, D, F) The percentage of the total volume represented by the expansion and acidified compartment volume. In panel B pie graphs, red and blue represent the percent of the expansion compartment and the acidified compartment, respectively. Each data point represents one lysosome; bars and whiskers represent the mean ± SEM. n=20-25 lysosomes from 3 worms (each of which is a biological replicate) for all graphs and all conditions (Table S1). One-Way Anova, ns, not significant, * P ≤ 0.05, ** P ≤ 0.01, *** P ≤ 0.001, **** P ≤ 0.0001.

**Supplemental Figure 2 (with main Figures 3 & 4).**
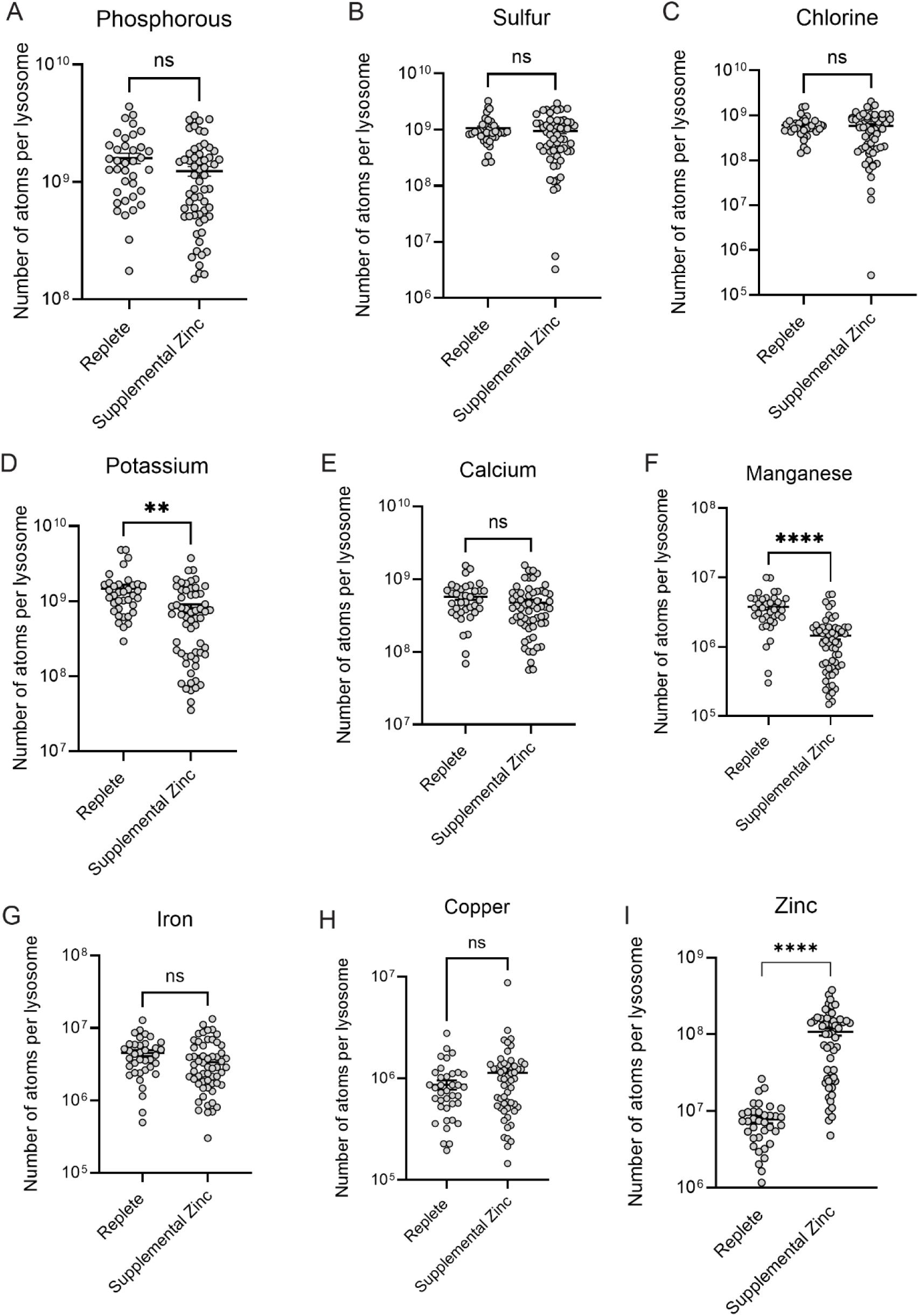
Comparison of elemental abundance in replete and zinc excess medium. (A-I) L4 stage transgenic worms expressing CDF::GFP and ZIPT-2.3::mCherry were transferred to NAMM dishes containing 0 or 200 μM supplemental zinc and cultured for 24 hours. Intestinal lysosomes were isolated by dissection one day after the L4 stage, and XFM Images were acquired with a 0.08μm step size and a dwell time of 500ms (N=34 lysosomes from 3 worms for replete; N=26 lysosomes from 3 worms for zinc excess). The number of atoms per lysosome for each element is displayed on a logarithmic scale. Each data point represents one lysosome; bars and whiskers represent the mean ± SEM. Comparison of the number of zinc atoms per lysosome from animals cultured in standard NAMM (replete) or 200 μM supplemental zinc (excess) was performed by Welch’s t test, ns, not significant, * P ≤ 0.05, ** P ≤ 0.01, *** P ≤ 0.001, **** P ≤ 0.0001. In excess zinc, potassium and manganese were significantly lower, zinc was significantly higher, and other elements were not significantly changed.

**Supplemental Figure 3 (with main Figure 5).**
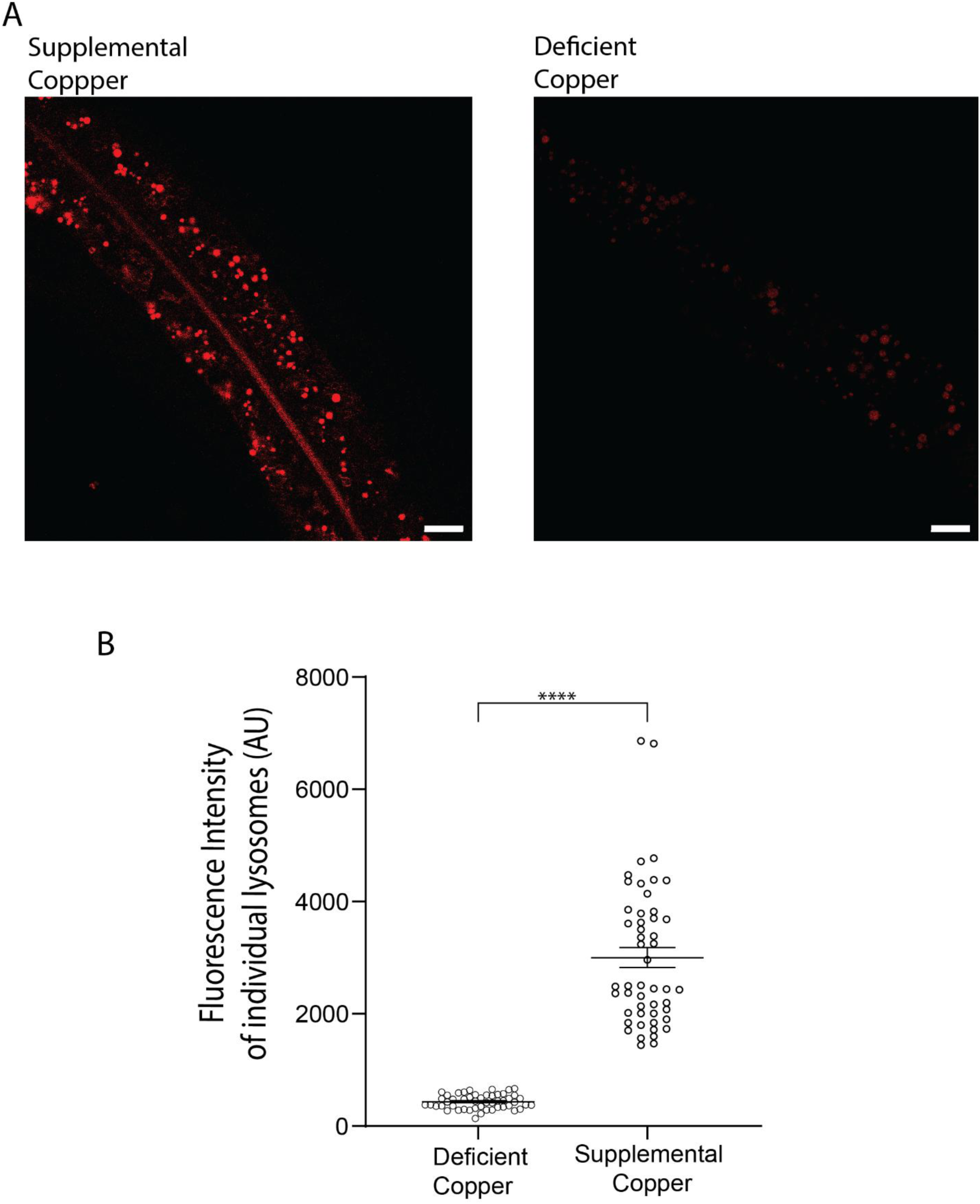
The CF4 dye detects copper in *C. elegans* intestinal lysosomes. (A) L4 stage wild-type worms were cultured for 24 hours on NAMM dishes with 25 μM CF4 dye and either 100μM supplemental copper or 50μM of BCS chelator (deficient). Worms were imaged by super-resolution microscopy for red fluorescence. Scale bar = 2μm. One representative image is shown for each condition. CF4 red fluorescence displays intense staining in intestinal lysosomes in copper excess, whereas it is only faintly visible in copper deficiency. (B) Fluorescence intensity of individual lysosomes in arbitrary units (AU) was measured by drawing a ROI around each lysosome in Fiji. Values for individual lysosomes are shown as circles; bar and whiskers represent the average +/- the standard error of the mean. For copper excess and deficiency, n=50 lysosomes from 3 animals. Welch’s t test, **** P ≤ 0.0001.

**Supplemental Figure 4 (with main Figure 5).**
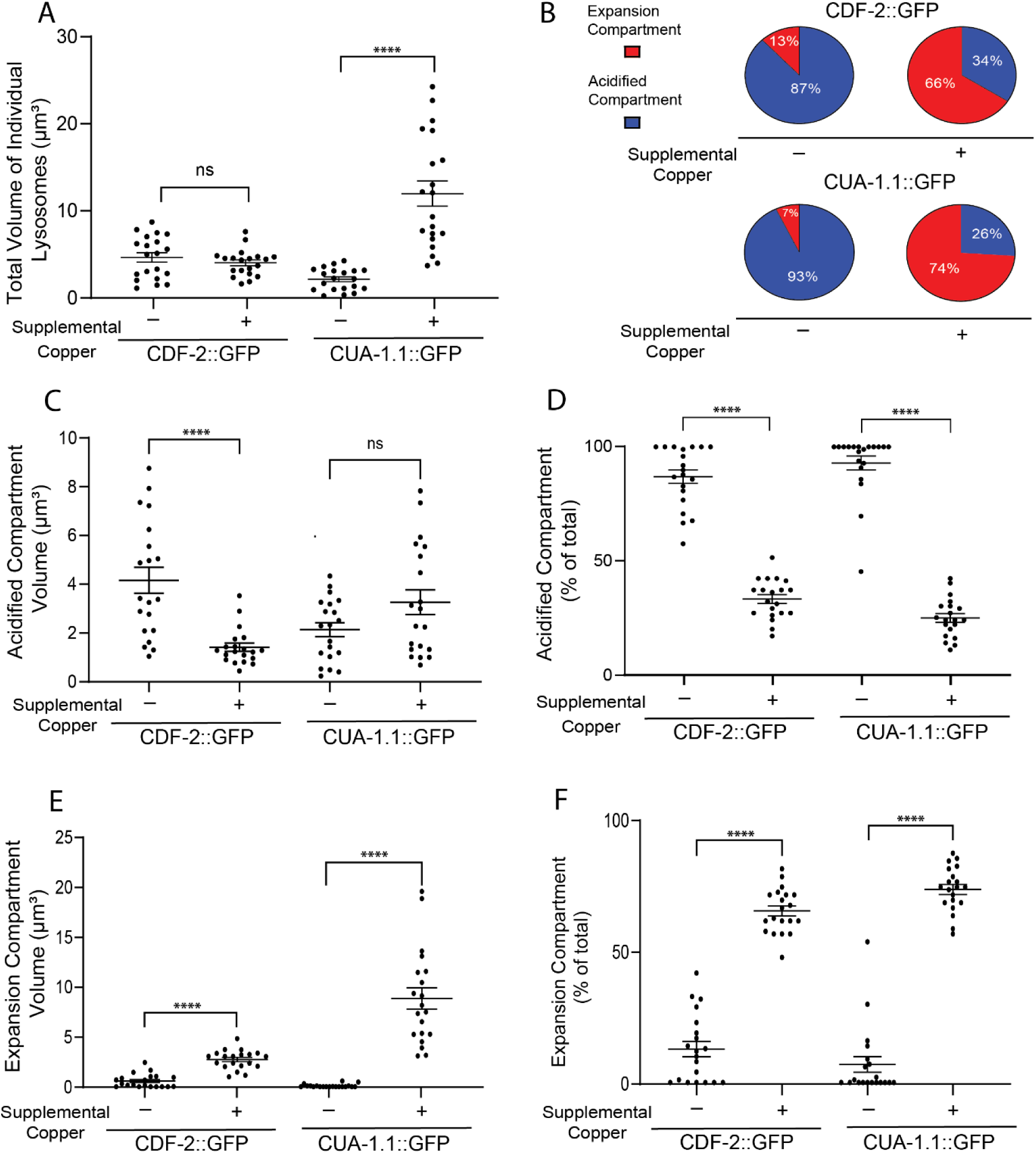
Quantification of acidified compartment volume, expansion compartment volume, and total lysosome volume. Transgenic L4 stage animals expressing CDF-2::GFP or CUA-1.1::GFP were cultured for 24 hours in 1 μM LysoTracker blue and 25 μM CF4 in NAMM medium (-supplemental copper) or NAMM medium containing 100μM copper (+ supplemental copper). Lysosomes were imaged at the young adult stage by super-resolution microscopy with green, red, and blue fluorescence. The volumes of the expansion compartment and the acidified compartment were calculated from 3-dimensional images by assuming the compartments are shaped like spheres (see Experimental Procedures for details). (A, C, E) Total lysosome, acidified compartment, and expansion compartment volume (μm_3_). (B, D, F) The percentage of the total volume represented by the expansion and acidified compartment volumes. In panel B pie graphs, red and blue represent the percent of the expansion compartment and the acidified compartment, respectively. Each data point represents one lysosome; bars and whiskers represent the mean ± SEM. n=20 lysosomes from 3 animals for all graphs and all conditions (Table S2). Statistical comparisons are between the same genotype in copper replete and copper excess. Welch’s t test: ns, not significant, * P ≤ 0.05, ** P ≤ 0.01, *** P ≤ 0.001, **** P ≤ 0.0001.

**Supplemental Figure 5 (with main Figure 6).**
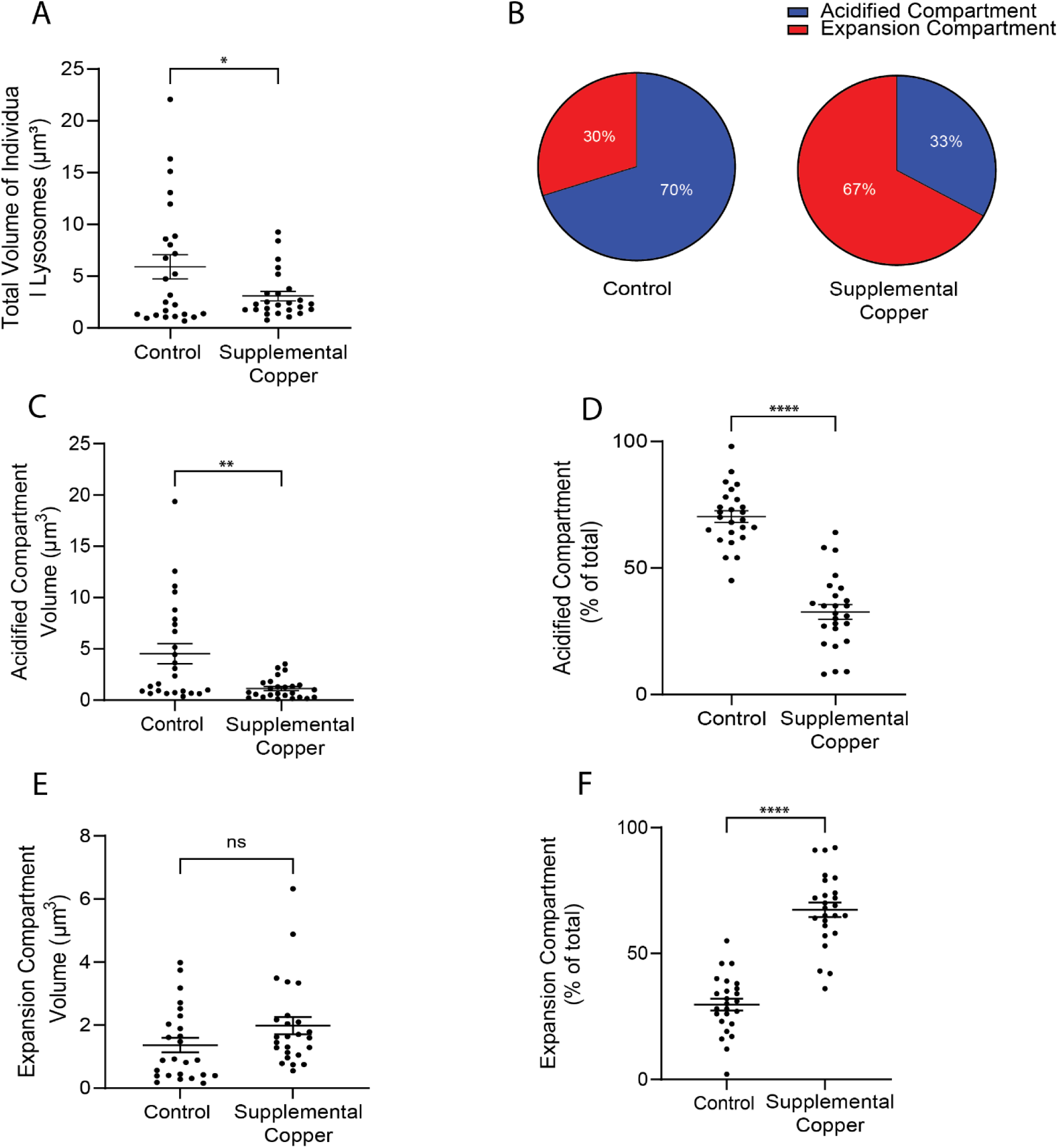
Quantification of acidified compartment volume, expansion compartment volume, and total lysosome volume. Transgenic L4 stage animals expressing CDF-2::mCherry and CUA-1.1::GFP were cultured for 24 hours in 1 μM LysoTracker blue and 25 μM CF4 in NAMM medium (control) or NAMM medium containing 100μM copper (supplemental copper). Lysosomes were imaged at the young adult stage by super-resolution microscopy with green, red, and blue fluorescence. The volumes of the expansion compartment and the acidified compartment were calculated from 3-dimensional images by assuming the compartments are shaped like spheres (see Experimental Procedures for details). (A, C, E) Total lysosome, acidified compartment, and expansion compartment volume (μm_3_). (B, D, F) The percentage of the total volume represented by the expansion and acidified compartment volumes. In panel B pie graphs, red and blue represent the percent of the expansion compartment and the acidified compartment, respectively. Each data point represents one lysosome; bars and whiskers represent the mean ± SEM. n=25 lysosomes from 3 animals for all graphs and all conditions (Table S3). Statistical comparisons are between the same genotype in copper replete and copper excess. Welch’s t test: ns, not significant, * P ≤ 0.05, ** P ≤ 0.01, *** P ≤ 0.001, **** P ≤ 0.0001.

**Table S1.**

| Control |  |  |  |  |  |
| --- | --- | --- | --- | --- | --- |
| Lysosome # | Total Volume ($\mu\text{m}^3$) | Acidified Compartment Volume ($\mu\text{m}^3$) | Expansion Compartment Volume ($\mu\text{m}^3$) | Acidified Compartment (% of total) | Expansion Compartment (% of total) |
| 1 | 1.37 | 0.988 | 0.380 | 72 | 28 |
| 2 | 3.24 | 2.81 | 0.435 | 87 | 13 |
| 3 | 1.55 | 0.608 | 0.938 | 39 | 61 |
| 4 | 1.72 | 1.14 | 0.586 | 66 | 34 |
| 5 | 1.51 | 1.12 | 0.397 | 74 | 26 |
| 6 | 3.57 | 3.57 | 0.000 | 100 | 0 |
| 7 | 2.02 | 1.16 | 0.855 | 58 | 42 |
| 8 | 1.38 | 0.505 | 0.876 | 37 | 63 |
| 9 | 2.12 | 0.798 | 1.32 | 38 | 62 |
| 10 | 5.14 | 4.04 | 1.09 | 79 | 21 |
| 11 | 1.54 | 0.887 | 0.649 | 58 | 42 |
| 12 | 0.43 | 0.406 | 0.022 | 95 | 5 |
| 13 | 2.85 | 1.09 | 1.76 | 38 | 62 |
| 14 | 1.43 | 0.885 | 0.540 | 62 | 38 |
| 15 | 2.93 | 1.77 | 1.16 | 60 | 40 |
| 16 | 1.89 | 0.854 | 1.03 | 45 | 55 |
| 17 | 2.00 | 0.448 | 1.56 | 22 | 78 |
| 18 | 1.59 | 0.549 | 1.04 | 34 | 66 |
| 19 | 1.76 | 0.925 | 0.831 | 53 | 47 |
| 20 | 3.28 | 2.29 | 0.995 | 70 | 30 |
| 21 | 15.1 | 15.1 | 0.000 | 100 | 0 |
| 22 | 14.6 | 14.6 | 0.000 | 100 | 0 |
| 23 | 14.9 | 13.9 | 0.982 | 93 | 7 |
| 24 | 14.4 | 14.4 | 0.00 | 100 | 0 |
| 25 | 5.58 | 4.83 | 0.752 | 87 | 13 |
| Averages | 4.31 | 3.59 | 0.728 | 67 | 33 |
| Zinc |  |  |  |  |  |
| Lysosome # | Total Volume ($\mu\text{m}^3$) | Acidified Compartment Volume ($\mu\text{m}^3$) | Expansion Compartment Volume ($\mu\text{m}^3$) | Acidified Compartment (% of total) | Expansion Compartment (% of total) |
| 1 | 20.5 | 3.35 | 17.1 | 16 | 84 |
| 2 | 19.2 | 4.29 | 14.9 | 22 | 78 |
| 3 | 19.9 | 3.90 | 16.0 | 20 | 80 |
| 4 | 6.40 | 1.57 | 4.83 | 25 | 75 |
| 5 | 2.49 | 0.606 | 1.89 | 24 | 76 |
| 6 | 9.66 | 1.09 | 8.57 | 11 | 89 |
| 7 | 12.4 | 2.17 | 10.2 | 18 | 82 |

| 8 | 24.4 | 3.78 | 20.6 | 15 | 85 |
| --- | --- | --- | --- | --- | --- |
| 9 | 10.8 | 3.54 | 7.26 | 33 | 67 |
| 10 | 26.4 | 6.42 | 20.0 | 24 | 76 |
| 11 | 7.84 | 1.38 | 6.46 | 18 | 82 |
| 12 | 14.6 | 2.32 | 12.3 | 16 | 84 |
| 13 | 9.18 | 1.25 | 7.93 | 14 | 86 |
| 14 | 17.5 | 1.91 | 15.6 | 11 | 89 |
| 15 | 14.8 | 0.840 | 13.9 | 6 | 94 |
| 16 | 21.3 | 3.33 | 18.0 | 16 | 84 |
| 17 | 6.66 | 0.73 | 5.93 | 11 | 89 |
| 18 | 9.06 | 2.41 | 6.65 | 27 | 73 |
| 19 | 18.5 | 3.71 | 14.8 | 20 | 80 |
| 20 | 41.4 | 2.48 | 38.9 | 6 | 94 |
| 21 | 44.3 | 3.09 | 41.2 | 7 | 93 |
| 22 | 42.8 | 2.96 | 39.8 | 7 | 93 |
| Averages | 18.2 | 2.60 | 15.6 | 17 | 83 |
| Copper |  |  |  |  |  |
| Lysosome # | Total Volume ($\mu\text{m}^3$) | Acidified Compartment Volume ($\mu\text{m}^3$) | Expansion Compartment Volume ($\mu\text{m}^3$) | Acidified Compartment (% of total) | Expansion Compartment (% of total) |
| 1 | 7.00 | 0.154 | 6.85 | 2 | 98 |
| 2 | 9.25 | 3.49 | 5.76 | 38 | 62 |
| 3 | 17.0 | 0.728 | 16.2 | 4 | 96 |
| 4 | 11.0 | 1.35 | 9.68 | 12 | 88 |
| 5 | 13.3 | 0.453 | 12.8 | 3 | 97 |
| 6 | 10.9 | 1.35 | 9.53 | 12 | 88 |
| 7 | 5.81 | 1.38 | 4.43 | 24 | 76 |
| 8 | 7.51 | 0.665 | 6.84 | 9 | 91 |
| 9 | 5.53 | 0.343 | 5.19 | 6 | 94 |
| 10 | 6.57 | 3.28 | 3.30 | 50 | 50 |
| 11 | 4.70 | 4.19 | 0.510 | 89 | 11 |
| 12 | 7.38 | 2.57 | 4.81 | 35 | 65 |
| 13 | 3.94 | 0.983 | 2.96 | 25 | 75 |
| 14 | 4.10 | 2.12 | 1.98 | 52 | 48 |
| 15 | 6.98 | 1.58 | 5.40 | 23 | 77 |
| 16 | 7.90 | 1.77 | 6.13 | 22 | 78 |
| 17 | 3.32 | 1.04 | 2.28 | 31 | 69 |
| 18 | 3.19 | 2.74 | 0.45 | 86 | 14 |
| 19 | 2.75 | 1.88 | 0.87 | 68 | 32 |
| 20 | 2.33 | 0.388 | 1.95 | 17 | 83 |
| 21 | 49.4 | 7.77 | 41.6 | 16 | 84 |

| 22 | 45.1 | 3.32 | 41.8 | 7 | 93 |
| --- | --- | --- | --- | --- | --- |
| 23 | 21.3 | 3.28 | 18.0 | 15 | 85 |
| 24 | 38.1 | 5.16 | 32.9 | 14 | 86 |
| Averages | 12.3 | 2.17 | 10.1 | 27.5 | 72.5 |
| Manganese |  |  |  |  |  |
| Lysosome # | Total Volume ($\mu\text{m}^3$) | Acidified Compartment Volume ($\mu\text{m}^3$) | Expansion Compartment Volume ($\mu\text{m}^3$) | Acidified Compartment (% of total) | Expansion Compartment (% of total) |
| 1 | 15.6 | 7.32 | 8.27 | 47 | 53 |
| 2 | 9.88 | 5.69 | 4.18 | 58 | 42 |
| 3 | 7.66 | 3.85 | 3.80 | 50 | 50 |
| 4 | 19.1 | 5.06 | 14.0 | 26 | 74 |
| 5 | 15.1 | 7.36 | 7.73 | 49 | 51 |
| 6 | 15.7 | 4.37 | 11.3 | 28 | 72 |
| 7 | 6.69 | 5.68 | 1.01 | 85 | 15 |
| 8 | 11.7 | 7.55 | 4.13 | 65 | 35 |
| 9 | 5.36 | 5.36 | 0 | 100 | 0 |
| 10 | 10.2 | 7.82 | 2.38 | 77 | 23 |
| 11 | 16.4 | 9.68 | 6.76 | 59 | 41 |
| 12 | 10.1 | 6.30 | 3.77 | 63 | 37 |
| 13 | 3.42 | 1.39 | 2.03 | 41 | 59 |
| 14 | 6.73 | 6.73 | 0 | 100 | 0 |
| 15 | 14.9 | 14.0 | 0.913 | 94 | 6 |
| 16 | 13.4 | 5.75 | 7.69 | 43 | 57 |
| 17 | 7.17 | 3.52 | 3.66 | 49 | 51 |
| 18 | 6.82 | 4.10 | 2.71 | 60 | 40 |
| 19 | 6.89 | 3.15 | 3.73 | 46 | 54 |
| 20 | 7.13 | 3.04 | 4.09 | 43 | 57 |
| 21 | 39.8 | 8.71 | 31.1 | 22 | 78 |
| 22 | 21.5 | 6.55 | 15.0 | 30 | 70 |
| 23 | 17.7 | 1.28 | 16.4 | 7 | 93 |
| 24 | 9.60 | 2.74 | 6.86 | 29 | 71 |
| 25 | 31.0 | 0.552 | 30.5 | 2 | 98 |
| Averages | 13.2 | 5.50 | 7.68 | 51 | 49 |
| Cadmium |  |  |  |  |  |
| Lysosome # | Total Volume ($\mu\text{m}^3$) | Acidified Compartment Volume ($\mu\text{m}^3$) | Expansion Compartment Volume ($\mu\text{m}^3$) | Acidified Compartment (% of total) | Expansion Compartment (% of total) |
| 1 | 1.86 | 0.351 | 1.51 | 19 | 81 |
| 2 | 1.12 | 0.532 | 0.591 | 47 | 53 |
| 3 | 4.07 | 2.91 | 1.16 | 71 | 29 |
| 4 | 4.40 | 4.20 | 0.207 | 95 | 5 |
| 5 | 6.59 | 1.44 | 5.15 | 22 | 78 |
| 6 | 1.45 | 0.389 | 1.07 | 27 | 73 |
| 7 | 6.47 | 2.59 | 3.89 | 40 | 60 |
| 8 | 7.75 | 3.12 | 4.63 | 40 | 60 |
| 9 | 9.24 | 3.87 | 5.37 | 42 | 58 |
| 10 | 6.57 | 0.576 | 5.99 | 9 | 91 |
| 11 | 6.01 | 3.77 | 2.24 | 63 | 37 |
| 12 | 7.24 | 7.24 | 0.000 | 100 | 0 |
| 13 | 5.90 | 0.399 | 5.50 | 7 | 93 |
| 14 | 12.5 | 1.95 | 10.6 | 16 | 84 |
| 15 | 8.93 | 1.91 | 7.01 | 21 | 79 |
| 16 | 12.2 | 3.01 | 9.24 | 25 | 75 |
| 17 | 8.11 | 2.51 | 5.60 | 31 | 69 |
| 18 | 7.03 | 1.93 | 5.10 | 27 | 73 |
| 19 | 4.66 | 1.54 | 3.12 | 33 | 67 |
| 20 | 8.65 | 2.55 | 6.10 | 29 | 71 |
| Averages | 6.54 | 2.34 | 4.20 | 38 | 62 |

**Table S2.**
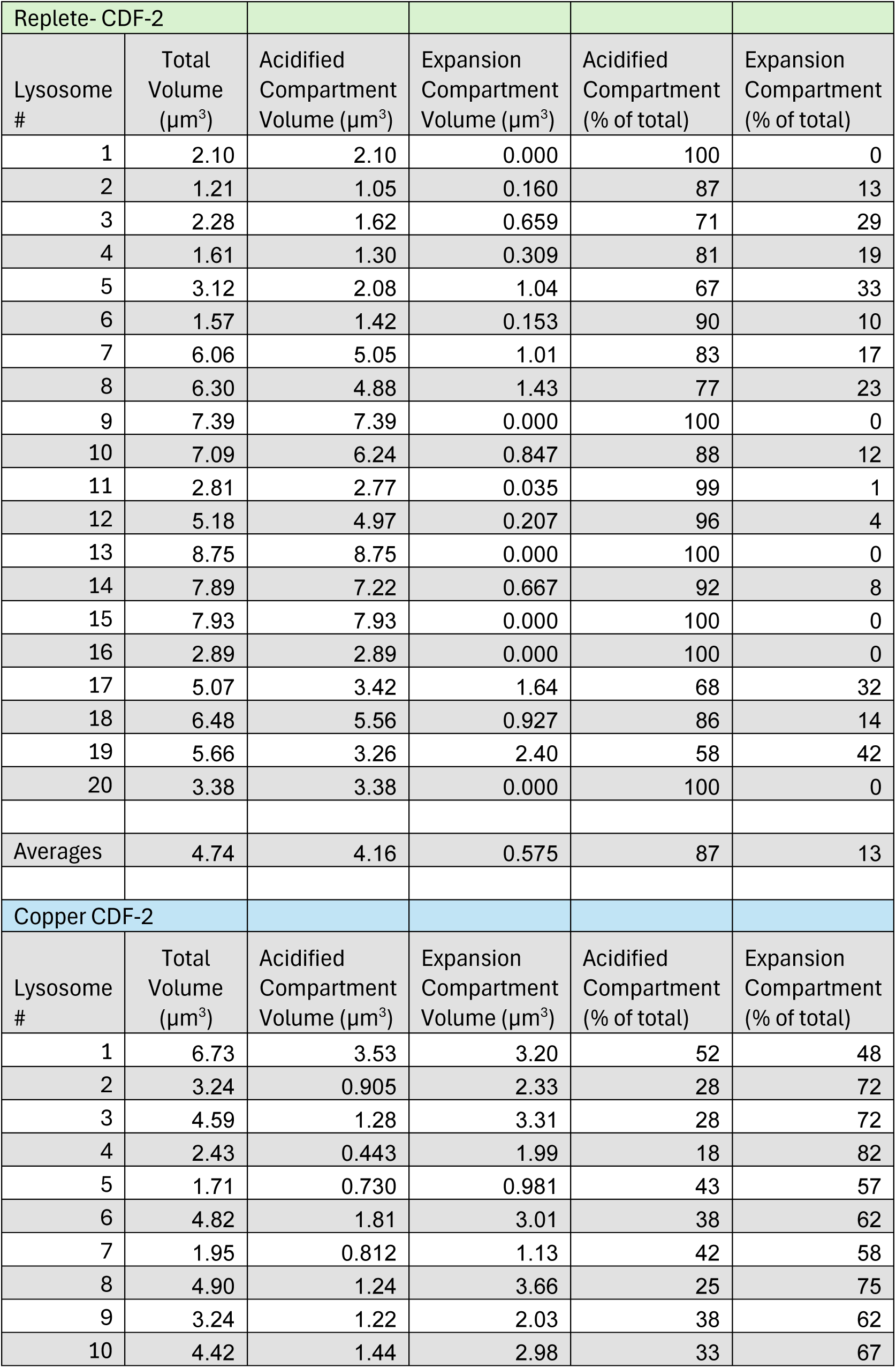
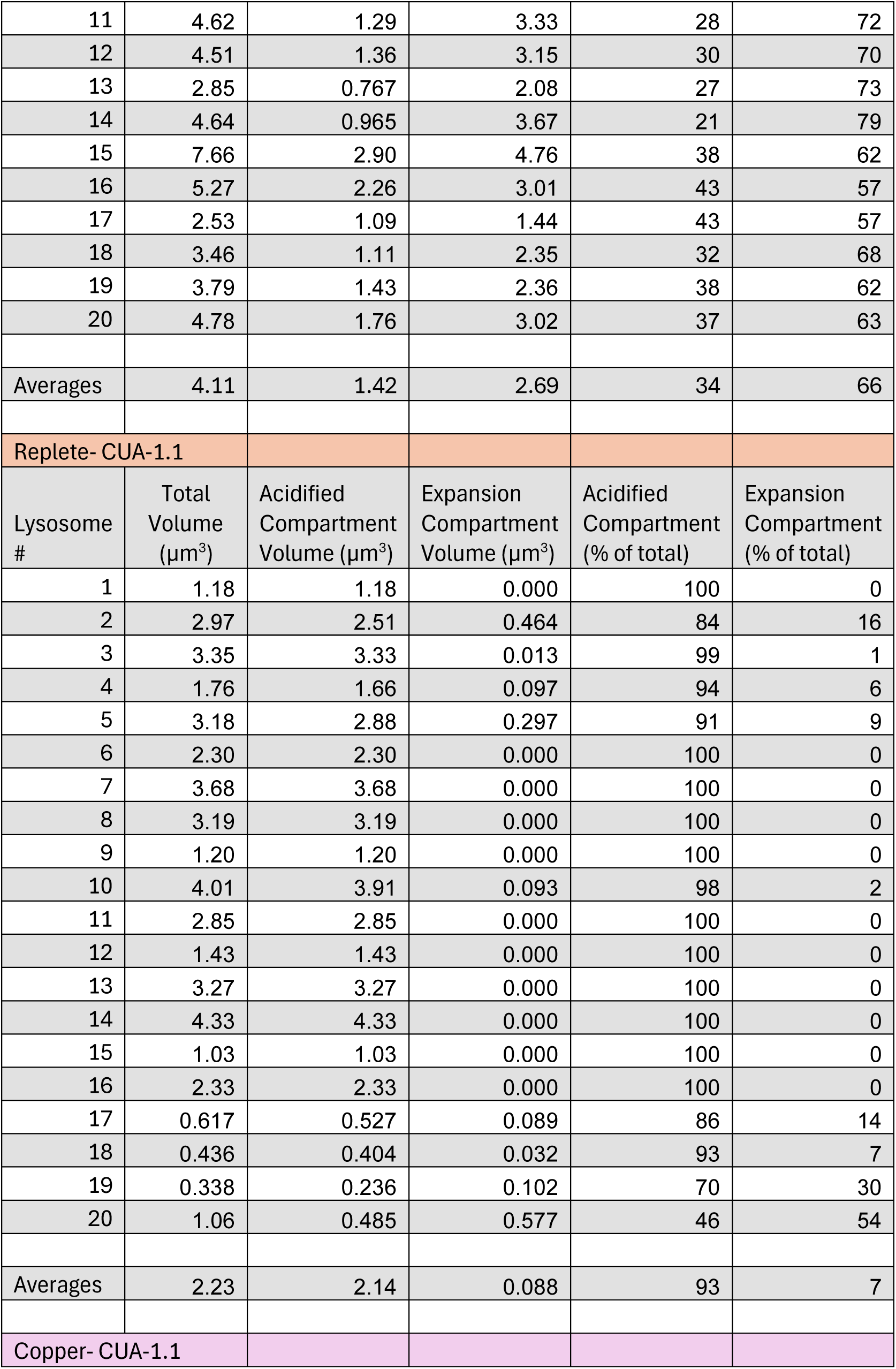
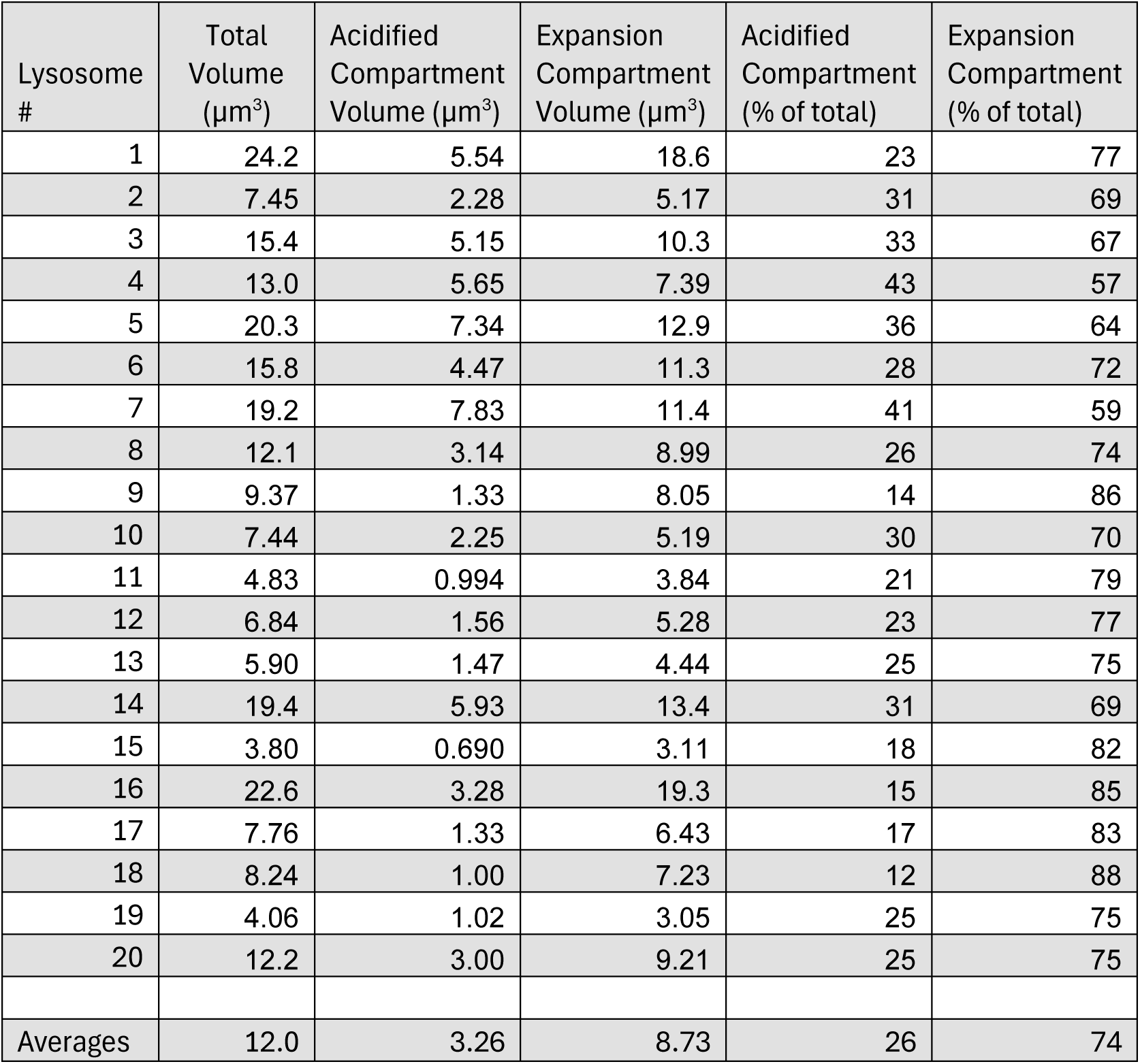

| Replete- CDF-2 |  |  |  |  |  |
| --- | --- | --- | --- | --- | --- |
| Lysosome # | Total Volume ($\mu\text{m}^3$) | Acidified Compartment Volume ($\mu\text{m}^3$) | Expansion Compartment Volume ($\mu\text{m}^3$) | Acidified Compartment (% of total) | Expansion Compartment (% of total) |
| 1 | 2.10 | 2.10 | 0.000 | 100 | 0 |
| 2 | 1.21 | 1.05 | 0.160 | 87 | 13 |
| 3 | 2.28 | 1.62 | 0.659 | 71 | 29 |
| 4 | 1.61 | 1.30 | 0.309 | 81 | 19 |
| 5 | 3.12 | 2.08 | 1.04 | 67 | 33 |
| 6 | 1.57 | 1.42 | 0.153 | 90 | 10 |
| 7 | 6.06 | 5.05 | 1.01 | 83 | 17 |
| 8 | 6.30 | 4.88 | 1.43 | 77 | 23 |
| 9 | 7.39 | 7.39 | 0.000 | 100 | 0 |
| 10 | 7.09 | 6.24 | 0.847 | 88 | 12 |
| 11 | 2.81 | 2.77 | 0.035 | 99 | 1 |
| 12 | 5.18 | 4.97 | 0.207 | 96 | 4 |
| 13 | 8.75 | 8.75 | 0.000 | 100 | 0 |
| 14 | 7.89 | 7.22 | 0.667 | 92 | 8 |
| 15 | 7.93 | 7.93 | 0.000 | 100 | 0 |
| 16 | 2.89 | 2.89 | 0.000 | 100 | 0 |
| 17 | 5.07 | 3.42 | 1.64 | 68 | 32 |
| 18 | 6.48 | 5.56 | 0.927 | 86 | 14 |
| 19 | 5.66 | 3.26 | 2.40 | 58 | 42 |
| 20 | 3.38 | 3.38 | 0.000 | 100 | 0 |
| Averages | 4.74 | 4.16 | 0.575 | 87 | 13 |
| Copper CDF-2 |  |  |  |  |  |
| Lysosome # | Total Volume ($\mu\text{m}^3$) | Acidified Compartment Volume ($\mu\text{m}^3$) | Expansion Compartment Volume ($\mu\text{m}^3$) | Acidified Compartment (% of total) | Expansion Compartment (% of total) |
| 1 | 6.73 | 3.53 | 3.20 | 52 | 48 |
| 2 | 3.24 | 0.905 | 2.33 | 28 | 72 |
| 3 | 4.59 | 1.28 | 3.31 | 28 | 72 |
| 4 | 2.43 | 0.443 | 1.99 | 18 | 82 |
| 5 | 1.71 | 0.730 | 0.981 | 43 | 57 |
| 6 | 4.82 | 1.81 | 3.01 | 38 | 62 |
| 7 | 1.95 | 0.812 | 1.13 | 42 | 58 |
| 8 | 4.90 | 1.24 | 3.66 | 25 | 75 |
| 9 | 3.24 | 1.22 | 2.03 | 38 | 62 |
| 10 | 4.42 | 1.44 | 2.98 | 33 | 67 |

| 11 | 4.62 | 1.29 | 3.33 | 28 | 72 |
| --- | --- | --- | --- | --- | --- |
| 12 | 4.51 | 1.36 | 3.15 | 30 | 70 |
| 13 | 2.85 | 0.767 | 2.08 | 27 | 73 |
| 14 | 4.64 | 0.965 | 3.67 | 21 | 79 |
| 15 | 7.66 | 2.90 | 4.76 | 38 | 62 |
| 16 | 5.27 | 2.26 | 3.01 | 43 | 57 |
| 17 | 2.53 | 1.09 | 1.44 | 43 | 57 |
| 18 | 3.46 | 1.11 | 2.35 | 32 | 68 |
| 19 | 3.79 | 1.43 | 2.36 | 38 | 62 |
| 20 | 4.78 | 1.76 | 3.02 | 37 | 63 |
| Averages | 4.11 | 1.42 | 2.69 | 34 | 66 |
| Replete- CUA-1.1 |  |  |  |  |  |
| Lysosome # | Total Volume ($\mu\text{m}^3$) | Acidified Compartment Volume ($\mu\text{m}^3$) | Expansion Compartment Volume ($\mu\text{m}^3$) | Acidified Compartment (% of total) | Expansion Compartment (% of total) |
| 1 | 1.18 | 1.18 | 0.000 | 100 | 0 |
| 2 | 2.97 | 2.51 | 0.464 | 84 | 16 |
| 3 | 3.35 | 3.33 | 0.013 | 99 | 1 |
| 4 | 1.76 | 1.66 | 0.097 | 94 | 6 |
| 5 | 3.18 | 2.88 | 0.297 | 91 | 9 |
| 6 | 2.30 | 2.30 | 0.000 | 100 | 0 |
| 7 | 3.68 | 3.68 | 0.000 | 100 | 0 |
| 8 | 3.19 | 3.19 | 0.000 | 100 | 0 |
| 9 | 1.20 | 1.20 | 0.000 | 100 | 0 |
| 10 | 4.01 | 3.91 | 0.093 | 98 | 2 |
| 11 | 2.85 | 2.85 | 0.000 | 100 | 0 |
| 12 | 1.43 | 1.43 | 0.000 | 100 | 0 |
| 13 | 3.27 | 3.27 | 0.000 | 100 | 0 |
| 14 | 4.33 | 4.33 | 0.000 | 100 | 0 |
| 15 | 1.03 | 1.03 | 0.000 | 100 | 0 |
| 16 | 2.33 | 2.33 | 0.000 | 100 | 0 |
| 17 | 0.617 | 0.527 | 0.089 | 86 | 14 |
| 18 | 0.436 | 0.404 | 0.032 | 93 | 7 |
| 19 | 0.338 | 0.236 | 0.102 | 70 | 30 |
| 20 | 1.06 | 0.485 | 0.577 | 46 | 54 |
| Averages | 2.23 | 2.14 | 0.088 | 93 | 7 |
| Copper- CUA-1.1 |  |  |  |  |  |

| Lysosome # | Total Volume ($\mu\text{m}^3$) | Acidified Compartment Volume ($\mu\text{m}^3$) | Expansion Compartment Volume ($\mu\text{m}^3$) | Acidified Compartment (% of total) | Expansion Compartment (% of total) |
| --- | --- | --- | --- | --- | --- |
| 1 | 24.2 | 5.54 | 18.6 | 23 | 77 |
| 2 | 7.45 | 2.28 | 5.17 | 31 | 69 |
| 3 | 15.4 | 5.15 | 10.3 | 33 | 67 |
| 4 | 13.0 | 5.65 | 7.39 | 43 | 57 |
| 5 | 20.3 | 7.34 | 12.9 | 36 | 64 |
| 6 | 15.8 | 4.47 | 11.3 | 28 | 72 |
| 7 | 19.2 | 7.83 | 11.4 | 41 | 59 |
| 8 | 12.1 | 3.14 | 8.99 | 26 | 74 |
| 9 | 9.37 | 1.33 | 8.05 | 14 | 86 |
| 10 | 7.44 | 2.25 | 5.19 | 30 | 70 |
| 11 | 4.83 | 0.994 | 3.84 | 21 | 79 |
| 12 | 6.84 | 1.56 | 5.28 | 23 | 77 |
| 13 | 5.90 | 1.47 | 4.44 | 25 | 75 |
| 14 | 19.4 | 5.93 | 13.4 | 31 | 69 |
| 15 | 3.80 | 0.690 | 3.11 | 18 | 82 |
| 16 | 22.6 | 3.28 | 19.3 | 15 | 85 |
| 17 | 7.76 | 1.33 | 6.43 | 17 | 83 |
| 18 | 8.24 | 1.00 | 7.23 | 12 | 88 |
| 19 | 4.06 | 1.02 | 3.05 | 25 | 75 |
| 20 | 12.2 | 3.00 | 9.21 | 25 | 75 |
| Averages | 12.0 | 3.26 | 8.73 | 26 | 74 |

**Table S3.**

| Replete |  |  |  |  |  |
| --- | --- | --- | --- | --- | --- |
| Lysosome # | Total Volume ($\mu\text{m}^3$) | Acidified Compartment Volume ($\mu\text{m}^3$) | Expansion Compartment Volume ($\mu\text{m}^3$) | Acidified Compartment (% of total) | Expansion Compartment (% of total) |
| 1 | 0.948 | 0.664 | 0.284 | 70 | 30 |
| 2 | 0.674 | 0.361 | 0.313 | 54 | 46 |
| 3 | 1.05 | 0.655 | 0.394 | 62 | 38 |
| 4 | 1.33 | 0.880 | 0.446 | 66 | 34 |
| 5 | 1.67 | 0.744 | 0.926 | 45 | 55 |
| 6 | 1.11 | 0.925 | 0.180 | 84 | 16 |
| 7 | 1.23 | 0.669 | 0.563 | 54 | 46 |
| 8 | 1.04 | 0.640 | 0.404 | 61 | 39 |
| 9 | 1.32 | 0.891 | 0.426 | 68 | 32 |
| 10 | 1.38 | 0.993 | 0.390 | 72 | 28 |
| 11 | 13.1 | 10.6 | 2.52 | 81 | 19 |
| 12 | 8.04 | 7.89 | 0.154 | 98 | 2 |
| 13 | 8.86 | 7.38 | 1.48 | 83 | 17 |
| 14 | 22.1 | 19.4 | 2.71 | 88 | 12 |
| 15 | 8.59 | 6.70 | 1.89 | 78 | 22 |
| 16 | 7.18 | 5.15 | 2.03 | 72 | 28 |
| 17 | 5.22 | 3.64 | 1.61 | 69 | 31 |
| 18 | 6.73 | 4.44 | 2.29 | 66 | 34 |
| 19 | 15.1 | 11.1 | 3.98 | 74 | 26 |
| 20 | 2.48 | 1.60 | 0.885 | 64 | 36 |
| 21 | 2.23 | 1.34 | 0.893 | 60 | 40 |
| 22 | 4.74 | 3.09 | 1.65 | 65 | 35 |
| 23 | 3.18 | 2.36 | 0.817 | 74 | 26 |
| 24 | 12.0 | 8.80 | 3.18 | 73 | 27 |
| 25 | 16.3 | 12.6 | 3.75 | 77 | 23 |
| Averages | 5.90 | 4.54 | 1.37 | 70 | 30 |
| Copper |  |  |  |  |  |
| Lysosome # | Total Volume ($\mu\text{m}^3$) | Acidified Compartment Volume ($\mu\text{m}^3$) | Expansion Compartment Volume ($\mu\text{m}^3$) | Acidified Compartment (% of total) | Expansion Compartment (% of total) |
| 1 | 2.30 | 1.34 | 0.964 | 58 | 42 |
| 2 | 1.76 | 1.01 | 0.750 | 57 | 43 |
| 3 | 2.31 | 0.690 | 1.62 | 30 | 70 |
| 4 | 9.26 | 2.94 | 6.32 | 32 | 68 |
| 5 | 6.64 | 3.15 | 3.49 | 47 | 53 |
| 6 | 2.04 | 1.30 | 0.742 | 64 | 36 |
| 7 | 3.78 | 1.48 | 2.30 | 39 | 61 |
| 8 | 1.78 | 0.503 | 1.28 | 28 | 72 |
| 9 | 1.08 | 0.296 | 0.781 | 27 | 73 |
| 10 | 2.42 | 0.634 | 1.78 | 26 | 74 |
| 11 | 2.22 | 0.765 | 1.45 | 35 | 65 |
| 12 | 1.40 | 0.112 | 1.29 | 8 | 92 |
| 13 | 1.79 | 0.159 | 1.63 | 9 | 91 |
| 14 | 0.76 | 0.215 | 0.547 | 28 | 72 |
| 15 | 1.34 | 0.287 | 1.05 | 21 | 79 |
| 16 | 1.74 | 0.148 | 1.59 | 9 | 91 |
| 17 | 2.52 | 0.476 | 2.04 | 19 | 81 |
| 18 | 1.88 | 0.581 | 1.30 | 31 | 69 |
| 19 | 3.32 | 1.15 | 2.17 | 35 | 65 |
| 20 | 8.42 | 3.53 | 4.88 | 42 | 58 |
| 21 | 3.35 | 1.25 | 2.10 | 37 | 63 |
| 22 | 1.41 | 0.286 | 1.13 | 20 | 80 |
| 23 | 5.19 | 1.82 | 3.37 | 35 | 65 |
| 24 | 5.82 | 2.48 | 3.34 | 43 | 57 |
| 25 | 2.67 | 0.967 | 1.70 | 36 | 64 |
| Averages | 3.09 | 1.10 | 1.99 | 33 | 67 |

